# A workflow for the identification of oligomeric structures on tilted sample planes in Cryo-SMLM

**DOI:** 10.64898/2026.05.12.724524

**Authors:** Yakun Dong, Zheyi Yang, Magdalena C. Schneider, Otmar Scherzer, Gerhard J. Schütz

## Abstract

We introduce a workflow to identify oligomeric structures that are recorded with single-molecule localization microscopy (SMLM) under cryogenic conditions. Typically, these oligomers are assumed to consist of protomers arranged as equilateral two-dimensional polygons and every protomer is labeled with a dye molecule for visualization. Unlike previous work, we consider scenarios in which the sample plane has an unknown orientation relative to the focal plane. Our contribution is a high-precision plane-fitting algorithm to determine the sample plane, combined with geometrical transformations and two circle-fitting algorithms to identify the oligomeric structures. Our simulations on synthetic data demonstrate that the proposed workflow achieves high accuracy in estimating both the unknown tilted plane and the oligomer size.

## 1. Introduction

Super-resolution microscopy has advanced rapidly over the past decades, overcoming the diffraction limit of conventional light microscopy [23]. A prominent technique in this field is single-molecule localization microscopy (SMLM) [28], which relies on the precise detection and localization of individual fluorescent emitters. However, acquiring localizations in SMLM typically requires the collection of hundreds to thousands of frames over several minutes to hours, posing significant challenges for maintaining sample stability while preserving the ultrastructure [14]. Cryo-fixation [33] has proven effective in addressing these challenges and is fully compatible with SMLM, leading to cryo-SMLM [34, 13, 35, 11].

A further advantage of cryogenic conditions is that the fluorophore dipole orientations remain fixed over time scales of several hours [37]. This stability enables improved discrimination between localizations corresponding to different nearby molecules [25], significantly improving the quality of data analysis. However, fixed dipole orientations can also introduce challenges in determining fluorophore positions. In SMLM, two-dimensional fluorophore positions are typically estimated by fitting an isotropic two-dimensional Gaussian function, as fluorophore dipoles are assumed to be freely rotating at room temperature. This rotational freedom leads, on average, to an isotropic, Gaussian-like point-spread function (PSF). For instance, Rieger and Stallinga [21] propose an analytical formula for calculating fluorophore positions, which has been shown to be computationally efficient [29, 30]. However, in cryo-SMLM, where dipole orientations are fixed, this approach becomes inaccurate and can introduce significant localization bias [29, 30, 12]. To circumvent this limitation, Hinterer et al. [7] proposed to leverage astigmatic imaging; further using estimates of the fixed dipole orientations, they achieved unbiased and precise single-molecule localizations reaching the Cramér-Rao lower bound (CRLB).

In recent years, the development of cryo-SMLM has enabled geometric analyses of biological structures at nanometer resolution [36, 16, 17]. One important aspect is the determination of oligomer structures based on the localizations of the individual dye molecules attached to the protomers. For example, Schneider et al. [25] developed a workflow for estimating the side lengths of simulated oligomeric structures modeled as equilateral two-dimensional polygons. However, this approach relies on idealized geometric assumptions. As illustrated in Fig 1, the workflow assumes that the oligomers either lie on the *xy* plane (parallel to the focal plane) or on a sample plane tilted by a known angle *θ* around the *x*-axis. In realistic experimental scenarios, however, the orientation of the sample plane is not precisely known. Workflows and algorithms, which can deal with this general situation are studied in this paper.

**Fig 1.**
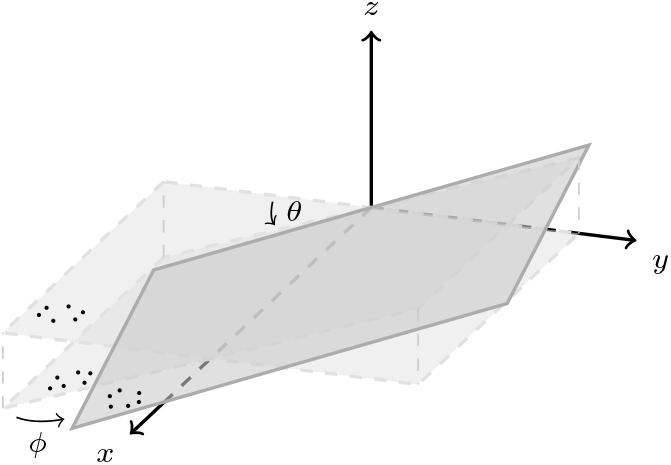
Illustration of the sample plane geometry. The z-axis represents the optical axis of the microscope. The xy-plane is the reference focal plane, which we call an ideal sample plane synonymously, meaning with zero tilt and zero shift. We assume that the protomers are perfectly lying in the sample plane. Any possible orientation of the sample plane is described by two parameters: the tilt angle θ around the x-axis and the rotation angle ϕ around the z-axis. To ensure the sample plane is positioned above the coverslip, an additional translation along the z-axis is applied (omitted in the figure for visual simplicity). This definition allows us to mathematically represent any sample plane in space.

The geometry of this sample plane can be parameterized by a tilt angle *θ* around the *x*-axis and a rotation angle *ϕ* around the *z*-axis, with *θ, ϕ≥* 0 (see Fig 1). For identification of the oligomeric structure on this unknown sample plane, we propose two workflows, the **Nominal workflow** and the **Robust workflow**. These workflows consist of five sequential steps (Table 4), differing mainly in the circle-fitting algorithm: (i) data preprocessing, (ii) plane-fitting (Algorithm 1) to estimate the sample plane orientation, (iii) coordinate transformations of the oligomer positions (Subsection 5.2.1), (iv) standard (Algorithm 2) or robust (Algorithm 3) circle-fitting method to estimate the circumcircle of the transformed oligomers as well as the side length of it, and (v) data postprocessing.

To distinguish our contributions from the previous work utilized in this manuscript, we illustrate the overall framework in Fig 2. Specifically, the key innovations of this paper can be summarized as follows:

**Fig 2.**
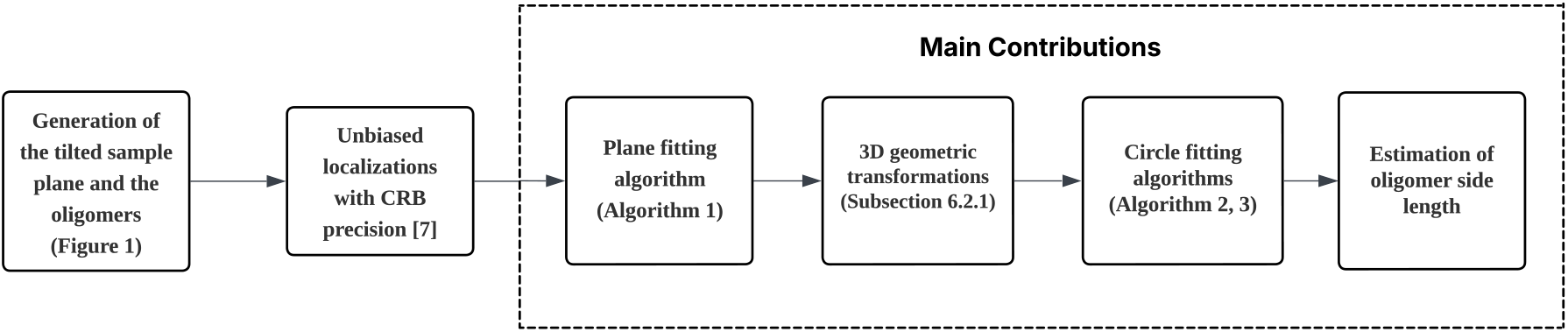
The proposed framework for identifying oligomeric structures on an unknown tilted sample plane. The diagram illustrates both the simulation steps generalized from prior work and the main contributions of this manuscript (dashed box).

1. **A unified algebraic framework for fast plane estimation and circle-fitting**. We utilize a non-iterative algebraic framework for both the unknown sample plane estimation (Algorithm 1) and the subsequent side length estimation via circle-fitting method (Algorithms 2, 3). By solving direct eigenvalue problems, this approach yields fast, high-precision estimators without requiring initial guesses. Furthermore, we extend this framework to address complex localization noise: the plane-fitting accommodates anisotropic noise, while the circle-fitting handles correlated noise.
2. **Problem-optimized transformations (Subsection 5.2.1)**. We propose a series of transformations that decouple the oligomer sizing procedure from the highly uncertain *z*-coordinates. By eliminating this dependence, we significantly reduce the overall structural estimation errors.

## 2. Results

In this section, we evaluate our proposed Nominal workflow and Robust workflow (summarized in Table 4) across four sets of simulations (Simulations I–IV, detailed in Table 1). Table 2 summarizes the main notations used throughout this chapter. These simulations explore various experimental settings, including small and large unknown tilt and rotation angles of the sample plane (Fig 1), varying maximum photon counts (Eq (4.1), with the detected photon counts *N*_eff_ defined in Eq (4.2)), and different oligomer configurations. Specifically within Simulation I, we also test a baseline case with a zero-tilt sample plane, which is consistent with the simulation setup used in [25]. The detailed simulation and evaluation steps are as follows. In each simulation, we generate 𝒩^oligo^ = 5000 well-separated oligomers with the same degree of oligomerization and side length on the sample plane, which is generalized from the code in [25]. For each oligomer *k* (*k* ∈{1, …, 𝒩^oligo^}), we number the protomers by index *i* (*i* ∈{1, …, *n*}, see Section 5). For each protomer, the corresponding localizations are evaluated using the software in [24]. We then assume that the localizations are accurately assigned based on the distinct dipole orientations of the fluorophore labels (see Subsection 4.5). To obtain an estimated position for each protomer, we average the corresponding localizations (Eq (5.2)). Using all estimated protomer positions within each simulation, we fit the tilted and rotated sample plane using our proposed algebraic plane-fitting method (Algorithm 1). Next, based on the problem-optimized transformations (Subsection 5.2.1), we estimate the original side length of the oligomer by applying our proposed circle-fitting methods (Algorithm 2 or Algorithm 3) to the oligomer positions after transformation. Depending on the chosen circle-fitting method, this process is referred to as the Nominal workflow or the Robust workflow. Finally, to obtain a representative side length for each simulation, we compute the mean, median, and weighted mean. The weighted mean estimator [3] is introduced specifically to downweight the side lengths obtained from oligomers with large localization errors. For each oligomer, we use the inverse of its post-transformation localization variance, i.e., 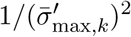, as a weight, where 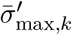 is defined in Eq (5.16). We now apply these established workflows and estimators to evaluate each simulation. The specific settings and results for each simulation are presented in the subsequent subsections.

**Table 1.** Ground-truth settings for the four sets of simulations (Simulations I–IV). The rows correspond to different simulation conditions: Simulation I varies the tilt and rotation angles; Simulation II varies the ground-truth side lengths; Simulation III varies the oligomerization state with a fixed radius; and Simulation IV varies the oligomerization state with a fixed side length. 𝒩^oliga^ = 5000 for all simulations.

| Simulation | Results | $\phi, \theta$ ($^\circ$) | $N_{\max}$ | Oligomerization | True side length $L^*$ (nm) |
| --- | --- | --- | --- | --- | --- |
| **I** | Fig 3 | 0, 0 | $5 \times 10^6$ | 4 | 5 |
|  |  | 10, 10 |  |  |  |
|  | Table 3 | 20, 20 |  |  |  |
|  |  | 30, 30 |  |  |  |
| **II** | Fig 4 | 10, 10<br>20, 20 | $5 \times 10^5, 5 \times 10^6$ | 4 | $\{1, 2, \dots, 10\}$ |
| **III** | Fig 5 | 10, 10<br>20, 20 | $5 \times 10^5, 5 \times 10^6$ | $n = \{3, 4, \dots, 8\}$ | $8 \sin\left(\frac{\pi}{n}\right)$ |
| **IV** | Fig 5 | 10, 10<br>20, 20 | $5 \times 10^5, 5 \times 10^6$ | $n = \{3, 4, \dots, 8\}$ | 5 |

**Table 2.** List of key symbols and their descriptions used in the Results section.

| Symbol | Description | Remarks |
| --- | --- | --- |
| $\mathcal{N}^{\text{oligo}}$ | Number of oligomers in one simulation | 5000 for each simulation |
| $n$ | Oligomerization, number of protomers in one oligomer | $n \in \{3, \dots, 8\}$ |
| $N_{\max}$ | Theoretical maximum number of detected photons (Eq 4.1) | $5 \times 10^5, 5 \times 10^6$ |
| $\theta, \phi$ | Tilt and rotation angle of the sample plane | See Fig 1 |
| $\hat{\theta}, \hat{\phi}$ | Estimated tilt, rotation of the sample plane | See Eq (5.12) |
| $L^*$ | True side length of the oligomer | few nanometers |
| $r^*$ | True circumscribed radius of the oligomer | $r^* = L^*/(2 \sin(\pi/n))$ |

### Simulation I: Varying tilt and rotation of the sample plane

We consider tilt and rotation angles of the sample plane increasing from *θ* = *ϕ* = 0° to 10°, 20°, and 30°, where the case of *θ* = 0° represents the zero tilt situation introduced in [25]. For this simulation set, we fix the maximum photon count at *N*_max_ = 5 ×10^6^, the oligomerization state at *n* = 4, and the ground-truth side length at *L*^*\**^ = 5 nm. We apply our proposed workflows for sizing oligomers and present the results in this part. Additionally, we compare our approach with the workflow in [25]. Since it is not directly applicable to our setting, we apply the necessary adaptations to enable the comparison. The detailed comparison results are therefore provided in the Supporting Information 7.3.

Fig 3 shows the histograms of the side lengths estimated by our proposed workflows. The peaks align with the ground-truth side length of 5 nm. We also summarize the results obtained by our workflows in Table 3, including the estimated sample plane and the mean, median, and weighted mean estimator of the reconstructed side lengths.

**Fig 3.**
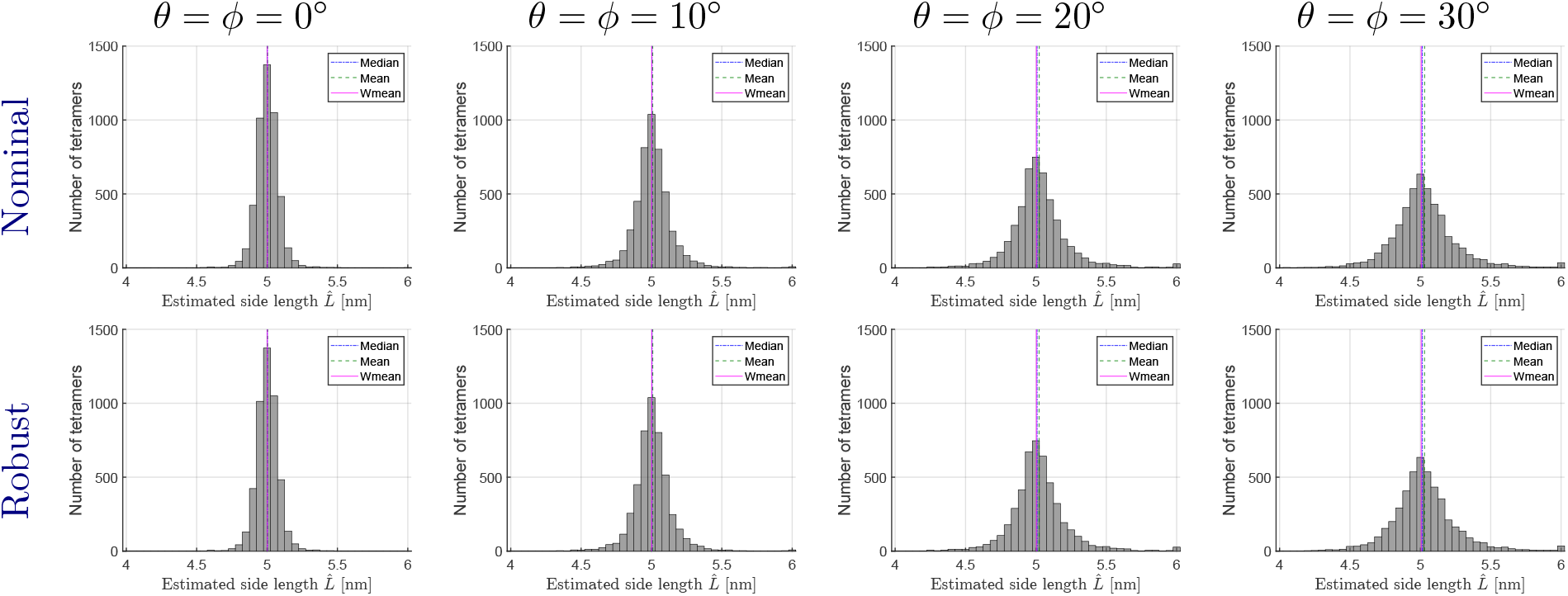
Histograms of estimated tetramer side lengths for Simulation I. The display range is 3.975 nm −6.025 nm (bin width: 0.05 nm), with outliers accumulated in the edge bins. The top and bottom rows correspond to the side length estimates by the Nominal workflow and the Robust workflow, respectively. Vertical lines indicate the weighted mean (solid magenta), mean (long-dashed green), and median (short-dashed blue) of the estimated side lengths for one simulation. Notably, the weighted mean estimator yields the most accurate results across all cases. Additionally, the Nominal workflow shows comparable performance to the Robust workflow, due to the precise localization enabled by the high photon count.

**Table 3.** Estimated data for Simulation I. The first column shows the ground-truth tilt and rotation of the simulated sample plane. Columns 3-6 show the tilt and rotation angles of the fitted plane, and the mean, median, and weighted mean of the estimated side length obtained using our two proposed workflows. Note that for 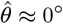, the fitted sample plane is fully characterized by the tilt angle 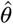alone, making the estimation of 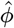 unidentifiable.

| $\theta/\phi$ ($^\circ$) | Workflow | $\hat{\theta}/\hat{\phi}$ ($^\circ$) | Mean (nm) | Median (nm) | W-Mean (nm) |
| --- | --- | --- | --- | --- | --- |
| 0/0 | Nominal workflow | 0.0001 / - | 5.0033 | 5.0021 | **5.0017** |
|  | Robust workflow |  | 5.0032 | 5.0021 | **5.0017** |
| 10/10 | Nominal workflow | 10.0000 / 10.0002 | 5.0088 | 5.0041 | **5.0024** |
|  | Robust workflow |  | 5.0090 | 5.0041 | **5.0024** |
| 20/20 | Nominal workflow | 20.0000 / 20.0001 | 5.0235 | 5.0087 | 5.0038 |
|  | Robust workflow |  | 5.0229 | 5.0087 | **5.0037** |
| 30/30 | Nominal workflow | 30.0000 / 30.0000 | 5.0307 | 5.0139 | 5.0065 |
|  | Robust workflow |  | 5.0300 | 5.0139 | **5.0064** |

**Table 4.** Summary of the two comprehensive workflows.

| Workflow | Description |
| --- | --- |
| **Nominal Workflow** | Data preprocessing (Section 4) → plane-fitting (Algorithm 1) → Geometric transformations (Subsection 5.2.1) → Algebraic circle-fitting (Algorithm 2) → Data postprocessing (Subsection 5.3). |
| **Robust Workflow** | Data preprocessing (Section 4) → plane-fitting (Algorithm 1) → Geometric transformations (Subsection 5.2.1) → Robust algebraic circle-fitting (Algorithm 3) → Data postprocessing (Subsection 5.3). |

As shown in the table, the plane-fitting (Algorithm 1) achieves high precision regardless of the tilt and rotation angles. However, larger tilt angles degrade the accuracy of the circle-fitting reconstruction, as shown in the histograms. Comparing the mean, median, and weighted mean estimates, we observe that the weighted mean is the best estimator for both of our proposed workflows. Furthermore, the Robust workflow shows slightly better performance than the Nominal workflow. This improvement is due to the robust circle-fitting algorithm’s downweighting of outliers (Algorithm 3). Particularly at critically low photon counts, the Robust workflow is much more accurate. For example, at *N*_max_ = 1 ×10^5^ (*θ* = *ϕ* = 10°), it estimates the side length at 5.0872 nm, whereas the Nominal workflow only achieves 5.0732 nm.

In the following simulations, we use *N*_max_ = 5×10^5^ and *N*_max_ = 5×10^6^ to represent cases with both reasonably larger and smaller localization errors. To improve the robustness and get a better estimation accuracy, we always employ the Robust workflow for the following estimates.

### Simulation II: Varying ground-truth side lengths

We consider varying the ground-truth side lengths of the oligomers *L*^*\**^ from 1 nm to 10 nm in this set of simulations. To compare the estimation precision, we first define the relative error of the reconstructed side length for the *k*-th oligomer as 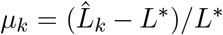, where *k* ∈ {1, …, 𝒩 ^oligo^}. We then compare the variability of the median relative error and the weighted mean relative error using 95 % bootstrap percentile confidence intervals estimated via a nonparametric bootstrap with 1000 resamples. We conduct this simulation under two tilt and rotation angles (*θ* = *ϕ* = 10° and 20°), and two different numbers of photons (*N*_max_ = 5 × 10^5^ and 5 × 10^6^).

Fig 4 shows the results estimated by the Robust workflow. The positive and negative values represent overestimation and underestimation, respectively. We observe that the weighted mean estimator still achieves lower relative errors than the median estimator, and estimates from simulations with a higher photon count, i.e., 5×10^6^, are more accurate than those with a lower photon count, 5×10^5^, due to the higher localization precision. As the ground-truth side length increases, we observe that the relative error of the estimated side length decreases, since a larger ground-truth side length reduces the impact of localization uncertainties.

**Fig 4.**
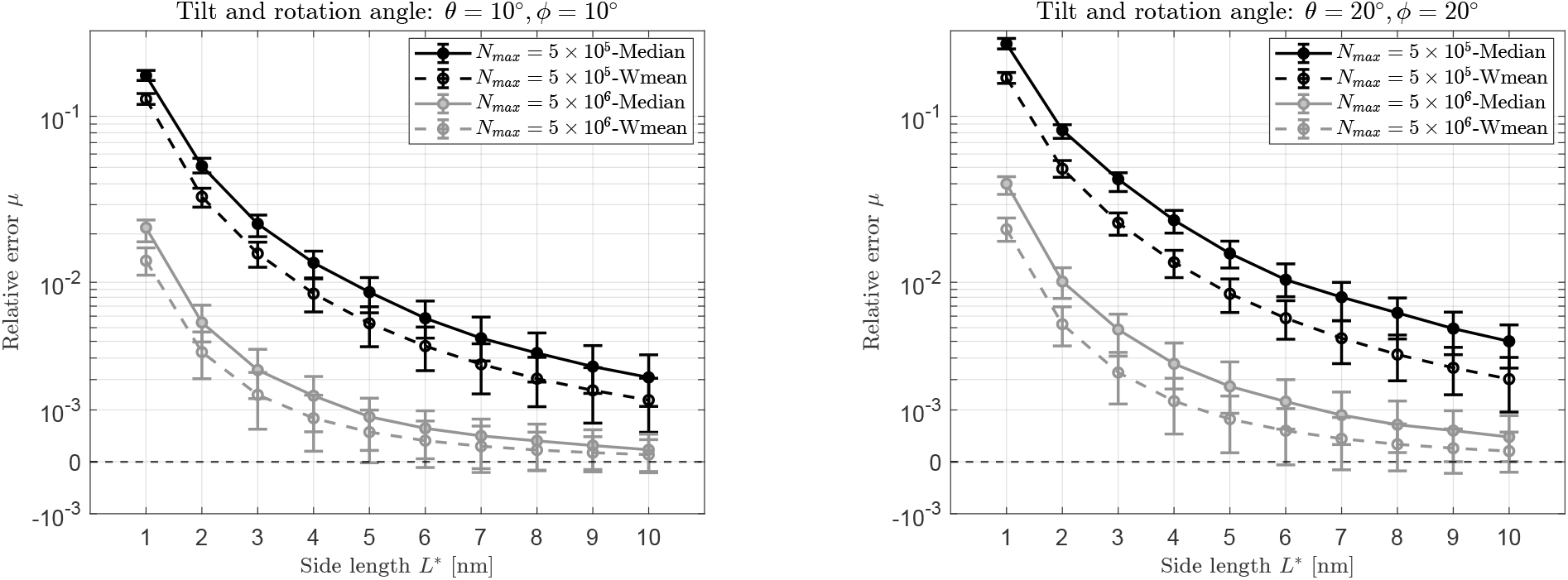
Relative error of reconstructed tetramer side lengths for varying ground-truth side lengths L^*^ ranging from 1 nm to 10 nm. The results indicate that estimation precision improves as the side length increases. As expected, the weighted mean estimator consistently yields the highest accuracy.

### Simulation III: Varying oligomerization with fixed radius

We vary the oligomerization *n* (*n* = 3, …, 8) with the circumscribed radius of the oligomer fixed at *r*^*\**^ = 4 nm. The ground-truth side length is thus given as 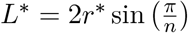. The simulations are carried out for two different photon numbers, *N*_*max*_ = 5 × 10^5^ and 5×10^6^, and two different tilt and rotation angles of the sample plane, *θ* = *ϕ* = 10° and *θ* = *ϕ* = 20°.

The estimates by the Robust workflow are shown in the first row of Fig 5. As the degree of oligomerization *n* increases, more vertices are available for fitting, which compensates for the potential instability caused by the decreasing side length. Consequently, the relative error exhibits a general downward trend, indicating that higher oligomerization states yield more accurate estimates.

**Fig 5.**
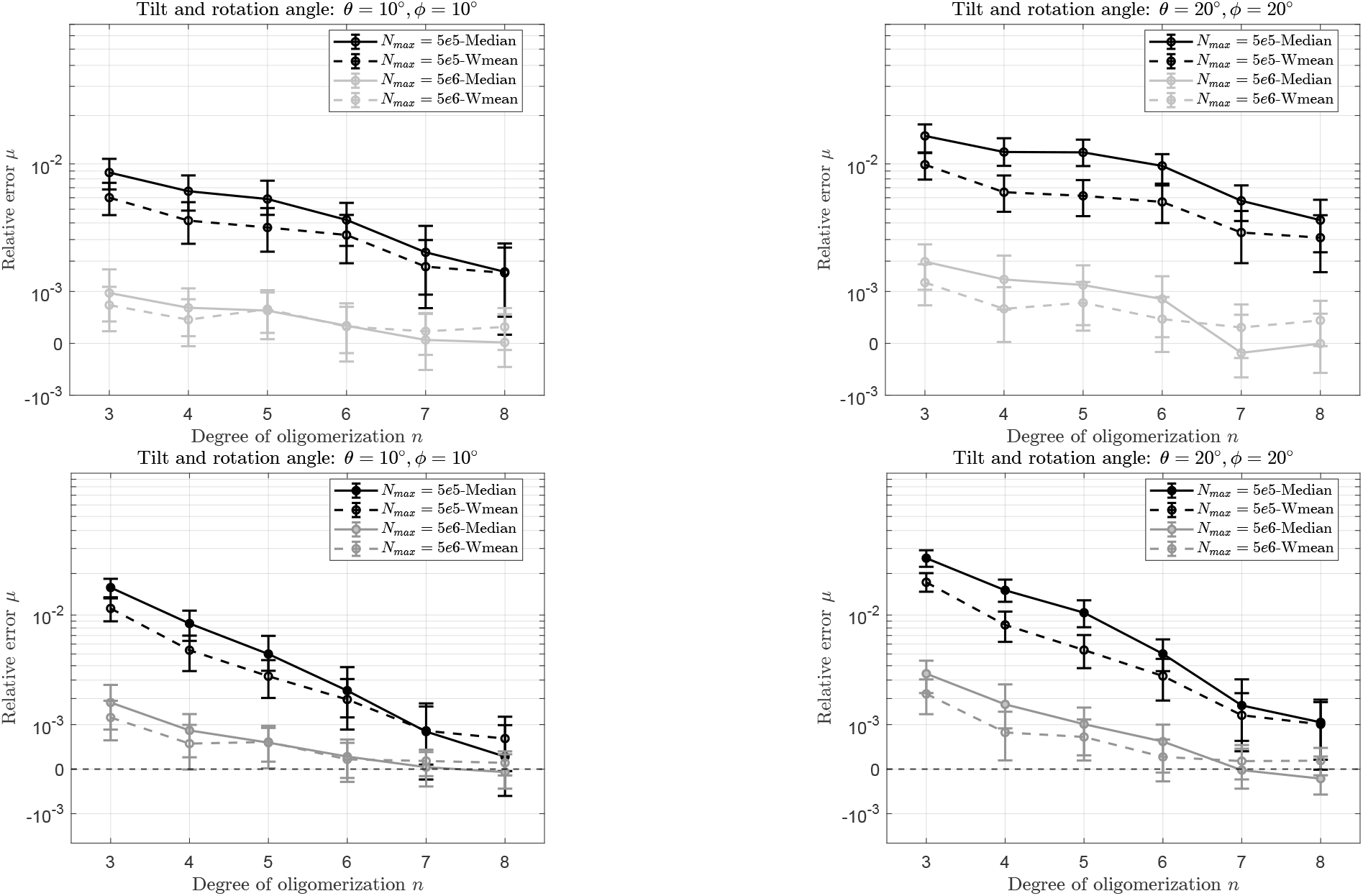
Relative error in side length estimation under fixed radius condition (1st row) or fixed side length condition (2nd row) by workflow 1.

### Simulation IV: Varying oligomerization with fixed side length

Similar to the Simulation III, we vary the oligomerization *n* (*n* = 3, …, 8), but with fixed side length of the oligomer *L*^*\**^ at 5 nm. The simulations are again carried out for two different photon numbers, *N*_max_ = 5 ×10^5^ and 5 10^6^, and two different tilt and rotation angles of the sample plane, *θ* = *ϕ* = 10° and *θ* = *ϕ* = 20°. The estimates obtained by the Robust workflow are shown in the second row of Fig 5. Since the side length is fixed in this case, the overall oligomer size increases with increasing oligomerization, thereby reducing the impact of localization errors on the side length estimation. Consequently, the estimates obtained here are more accurate than those from Simulation III.

## 3. Discussion

In practical cryo-SMLM experiments, the exact orientation of the sample plane is often unknown and potentially tilted. However, existing methods typically assume the sample plane is either perfectly aligned with the focal plane or precisely known, rendering them unrealistic for many applications. Our proposed workflows accurately resolve oligomeric structures under these general conditions, as demonstrated in Section 2. Building upon these results and our theoretical foundations (Sections 4 and 5), we highlight the following key points regarding the advantages and underlying assumptions of our framework:

i. **Generalized geometric modeling**. Unlike the restricted sample plane settings in previous work [25], our formulation of the sample plane can represent any plane in three-dimensional space (Fig 1), regardless of whether it is tilted or parallel relative to the focal plane. Importantly, we do not require prior knowledge of this sample plane; instead, it is directly estimated by our plane-fitting algorithm (Algorithm 1).
ii. **Generalization of the estimating method to account for anisotropic and correlated noise**. To achieve precise localization, we employ the unbiased estimator from [7], which inherently introduces anisotropic localization errors, complicating our estimation methods. To address this, we explicitly design our plane-fitting method to accommodate such noise and derive a high-precision sample plane estimation. For the circle-fitting, although the subsequent transformation step (Subsection 5.2.1) eliminates the impact of significant *z*-localization errors, it inevitably correlates the *x*- and *y*-localization errors. To overcome this, our circle-fitting algorithms (Algorithms 2 and 3) are generalized to operate under such correlated noise.
iii. **Practicality of the physical assumptions**. We assume that the degree of oligomerization can, in principle, be inferred from SMLM data by identifying the maximum number of localization clusters per oligomer. Additionally, localizations can be assigned to specific molecules by leveraging the measured dipole polarization. These physical priors allow us to effectively filter the oligomers, retaining only those with the correct number of localization clusters for subsequent estimations.

## 4. Data generation and Preprocessing

In this section, we describe the data generation and preprocessing steps used to prepare for the structural estimation. We first introduce the oligomer structure, sample plane orientation, fluorophore photophysics, and the simulation of localization errors. We then present the preprocessing steps, including the identification of oligomeric structures and the assignment of localizations to individual molecules. These procedures mainly follow the methodologies introduced in [25] and [7].

### 4.1. Sample plane and oligomer geometry

In the first step of our simulations, we utilize the source code provided in [25] to generate 𝒩^oligo^ oligomers on the reference *xy*-plane (which represents an ideal sample plane with zero tilt). These oligomers (*n*-mers, where *n*∈{3, 4, 5, 6, 7, 8}, corresponding to tri-through octamers) are modeled as regular polygons with fixed side lengths and random in-plane orientations. Subsequently, to simulate the actual orientation of the sample plane in this paper, we tilt and rotate this reference plane by *θ* and *ϕ* degrees, respectively, with *θ, ϕ ≥*0, as illustrated in Fig 1. Finally, an additional translation along the *z*-axis is applied to make the entire sample plane above the coverslip. Correspondingly, we set the focal plane of the simulated microscope to the center of the sample plane by assigning the nominal defocus value to the mean *z*-coordinate of all oligomers in one simulation.

### 4.2. Fluorophore photophysics and excitation model

At the molecular level, we assume that each protomer within the oligomer is labeled with exactly one fluorophore. To account for imaging under cryogenic conditions, a fixed dipole orientation is assigned to each fluorophore, which is independently and randomly generated for each molecule. The photophysical dynamics of these fluorophores incorporate realistic blinking characteristics [22], where the switching between bright and dark states follows a log-normal distribution in time [2].

During the imaging process, the fluorophores are alternately excited using differently polarized light. Since the absorption probability depends on the fluorophore’s dipole orientation, the detected numbers of emitted photons, *N*_*x*_ and *N*_*y*_, corresponding to excitation light polarized along the *x*- and *y*-axes respectively, are modeled as

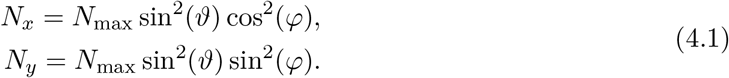

Here, *ϑ* and *φ* are the inclination and azimuth angles of the fluorophore’s dipole relative to the *z*- and *x*-axis, respectively (see Fig 6). *N*_max_ represents the maximum possible number of detected photons, achieved when the dipole is parallel to the polarization vector of the excitation light. Then the total number of detected photons *N*_eff_ is given as,

**Fig 6.**
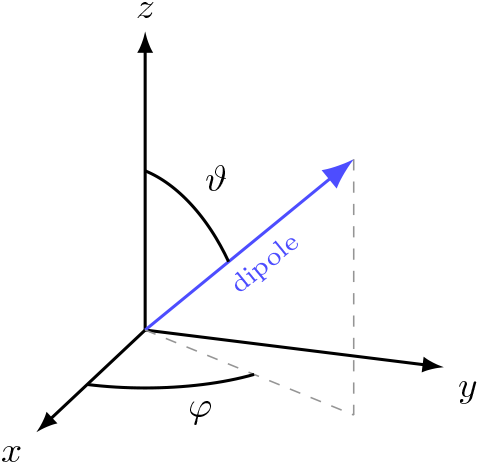
Illustrative dipole orientation

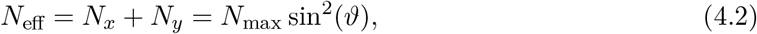

in which we implicitly apply Poisson shot noise to the detected number of photons to model signal fluctuations. In particular, for *ϑ* = 0, the dipole orientation is orthogonal to the excitation polarizations, resulting in no detected photons. In contrast, if *ϑ* = *π/*2, the maximum number of photons is detected.

### 4.3. Simulation of localizations

Following the methodology in [7], the ground-truth coordinates are first uniformly generated within the central pixel of a 17×17 ROI (108 nm/pixel) and a *±*500 nm axial range. We then calculate the Cramér-Rao bound using a vectorial PSF model [7, Eq 3], which is computationally implemented via the interactive simulation software developed by Schneider et al. [24]. As noted in Eq (4.2), this bound implicitly includes signal fluctuations due to Poisson shot noise. Finally, we simulate localizations of each protomer by adding these zero-mean, Cramér-Rao bound errors to the corresponding ground-truth coordinates. Due to the nature of the vectorial PSF model, the resulting localization errors are inherently anisotropic. This characteristic introduces specific difficulties for the plane-fitting analysis discussed in Section 5.

### 4.4. Identification of individual oligomeric structures

We assume that individual oligomers are well separated from one another, so that localizations can unambiguously be assigned to the correct oligomer based on spatial clustering.

### 4.5. Assignment of localizations to specific molecules

We assume that the oligomers are fully labeled, meaning each protomer is attached with exactly one fluorophore molecule. To correctly assign these localizations to their respective protomers, we utilize the measured dipole polarizations [35, 33]. Following a clustering preprocessing step analogous to [25], we retain only well-resolved oligomers that form exactly *n* distinct intensity clusters. For these oligomers, we further assume perfect localization assignment to individual molecules and use them for subsequent analysis.

## 5. Mathematical formulation and estimation algorithms

In this section, we define the mathematical models for the oligomeric structures and their localizations. We then introduce our plane-fitting method (Algorithm 1) to estimate the tilted sample plane. Subsequently, the coordinate transformations (Subsection 5.2.1) and two circle-fitting methods (Algorithms 2 and 3) are presented to determine the oligomer side lengths. Next, we provide two strategies for postprocessing the reconstructed side length (Subsection 5.3).

First, we establish the mathematical notations used for the subsequent analysis. We distinguish between oligomers, their protomers, and the associated localizations using the following indices: (i) Oligomers, *k* ∈ {1, …, 𝒩 ^oligo^}, representing the discrete complexes distributed on the sample plane; (ii) Protomers, *i* ∈ {1, …, *n*} within each oligomer, with ground-truth 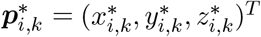 lying on the sample plane; and (iii) Localizations, *j* {1, …, *m*_*i,k*_} for each protomer, denoted by

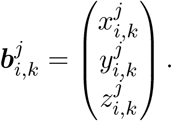

Since these localizations deviate from the ground-truth positions 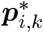 due to measurement uncertainties, this relation can be expressed as:

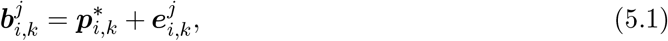

where 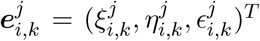 is an independent and identically distributed (i.i.d.) zero-mean Gaussian random vector. Specifically 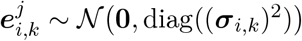, with the variance vector given (*σ*_*i,k*_) ^2^ = ((*σ*_*x,i,k*_) ^2^, (*σ*_*y,i,k*_) ^2^, (*σ*_*z,i,k*_)^2^)^*T*^.

As the localizations can be correctly assigned to their respective protomers (see Subsection4.5), we can obtain an estimate for the protomer position by computing the center of mass of the corresponding random localizations, defined as

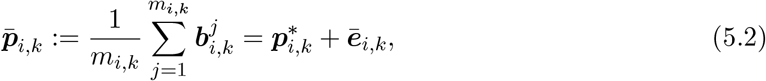

where the protomer position error 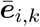is given by

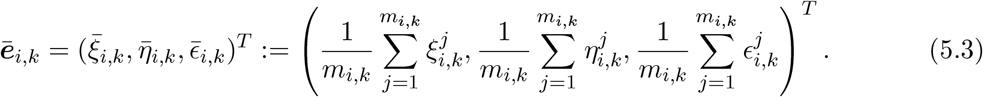

From which, we can easily verify that 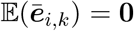, and the variance for 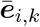, which is also the variance for protomer position, is given by,

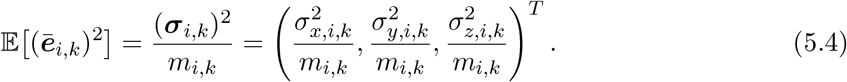

Consequently, 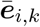follows a zero-mean normal distribution with independent components, denoted as 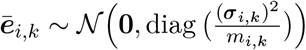.

Next, we connect our mathematical formulas with the localization simulations described in Subsection 4.3. From these simulations, the localization 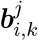 and its corresponding standard deviation 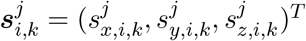 are provided as known inputs. Incorporating these given values, we will construct a variance estimator for protomer position 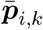.

Let 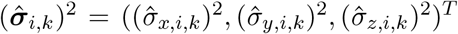denote the estimator for the localization variance (***σ***_*i,k*_)^2^. This estimator can be computed by averaging the given sample variances, i.e.,

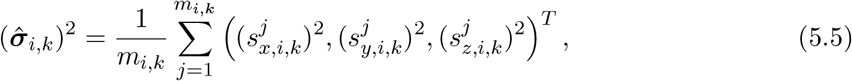

Ultimately, let 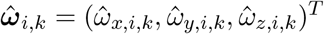denote the variance estimator for protomer position 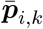,by substituting 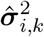 from Eq (5.5) into Eq (5.4), we derive 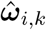as,

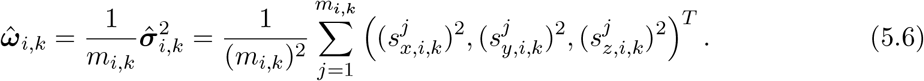

In the next subsection we will base on the estimated protomer positions 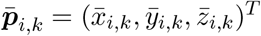and their corresponding position errors 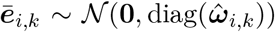to estimate the tilted sample plane.

### 5.1. Estimation of the tilted sample plane

We aim to estimate the parameter vector 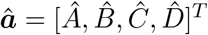 that defines the tilted sample plane 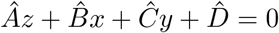. We can deriveâ by minimizing the mean squared algebraic error between the protomers and the plane. Defining the variable vector ***a*** = [*A, B, C, D*]^*T*^, this least square optimization is formulated as

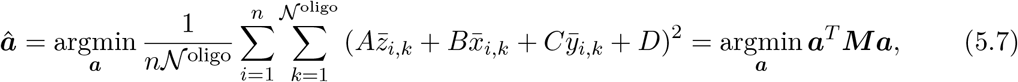

where ***M*** ∈ ℝ^4×4^ is defined as 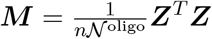, and the data matrix 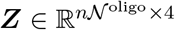 is given by

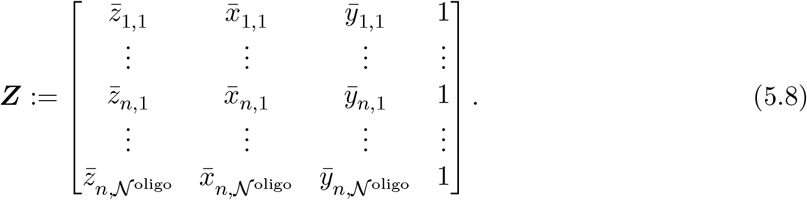

The typical values of *n* and 𝒩^oligo^ used in our simulations are summarized in Table 2. To avoid the trivial solution [0, 0, 0, 0]^*T*^ and derive a certain bias accuracy of the algebraic estimator, people usually impose a constraint ***a***^*T*^ ***Na*** = 1 on the minimization problem, where ***N***∈ℝ^4×4^ is the constraint matrix to be designed accordingly. By incorporating the constraint using a Lagrange multiplier *γ*, we formulate the min-max problem:

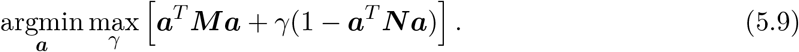

The optimal parameter vector ***â*** is obtained when ***a*** and *γ* satisfy the optimality conditions: ***Ma*** = *γ****Na*** and ***a***^*T*^ ***Na***− 1 = 0. By multiplying both sides of the first condition by ***a***^*T*^ and substituting the constraint, we derive *γ* = ***a***^*T*^ ***Ma***. Thus, the problem (5.9) is equivalent to finding the smallest positive generalized eigenvalue *γ* and its corresponding generalized eigenvector ***â*** by solving the equation

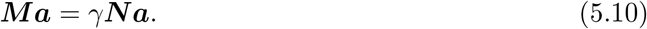

Similar to the error analysis routine introduced in [9, 27], we apply the matrix perturbation method [31] to Eq (5.10) to construct a constraint matrix ***N***. As demonstrated in the Supporting Information 7.1, this constraint matrix ensures that the plane parameter ***â*** remains unbiased up to the second order of the error.

Computationally, however, the constraint matrix ***N*** is typically not positive definite (which also holds true for the matrix provided in the Supporting Information 7.1), directly solving Eq (5.10) for the smallest positive generalized eigenvalue can be numerically unstable. To circumvent this, we reformulate it into the following equivalent generalized eigenvalue problem:

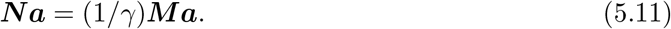

In the presence of noise, the data matrix ***M*** is strictly positive definite and *γ*≠ 0, ensuring the solvability of this problem. This allows us to find the generalized eigenvector ***â*** corresponding to the largest positive eigenvalue term (1*/γ*), which significantly improves the numerical stability of the algorithm. We then summarize the algebraic plane-fitting method in Algorithm 1.

#### Algorithm 1

Algebraic Plane-Fitting Method

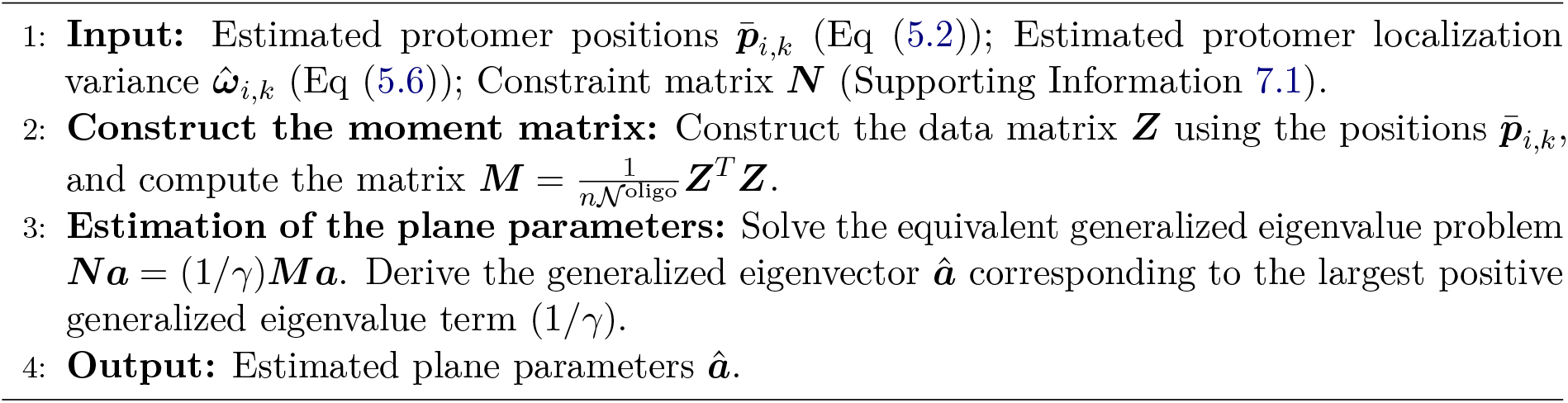

Ultimately, using the algebraic plane-fitting method described in Algorithm 1, we obtain the parameter vector 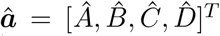 of the sample plane, which defines the plane equation 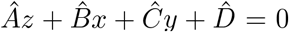. Subsequently, by analyzing the geometric orientation of the sample plane’s normal vector (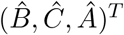, *Ĉ, Â*)^*T*^, the estimated tilt angle 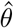 and rotation angle 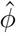 can be derived as follows:

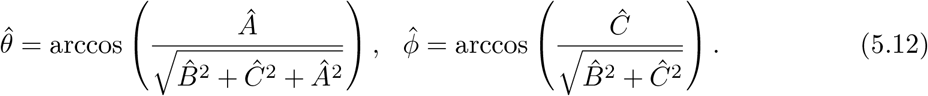

To simplify subsequent analysis, we treat the estimated 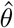 and 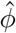 as fixed, deterministic real numbers.

### 5.2. Sizing the oligomers with circle-fitting algorithms

To estimate the size of the oligomer, we estimate its side length by calculating the radius of the circumscribed circle. For this calculation, we use a two-dimensional projection approach rather than estimating oligomer size directly on the tilted sample plane. This is because the localization error along the *z*-axis in SMLM is typically much larger than the errors in the *x* and *y* directions. To circumvent the big *z*-direction uncertainty in the circle-fitting procedure, we first project the estimated protomer positions 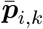 onto the *xy*-plane. However, this projection distorts the actual size and shape of the oligomers. To correct this geometric distortion, we apply a sequence of coordinate transformations to the projected points. As detailed in the following subsection, these transformations use the estimated angles 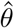 and 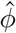 of the sample plane to restore the true geometry of the oligomer. This ensures that the circumscribed circle of the transformed two-dimensional points matches its true size on the sample plane, allowing us to compute the original oligomer’s side length.

#### 5.2.1 Transformations of the localizations and the localization errors

We begin by projecting the protomer positions 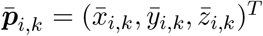 onto the *xy*-plane. The projected coordinates in the *xy*-plane can thus be written as 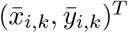, along with the projected localization error 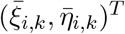. Next, we rotate the projected 2-dimensional coordinates and the associated localization errors by 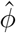 degrees around the *z*-axis, following the left-hand rule, and subsequently scale the *y*-coordinates of both rotated coordinates and rotated *y*-localization errors by the factor 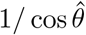, where 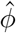 and 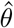 are the estimated rotation and tilt angle of the sample plane given in Eq (5.12). Using a simple, noise-free hexagon as an example, Fig 7 demonstrates that the proposed transformations can accurately restore the structure to its original size.

**Fig 7.**
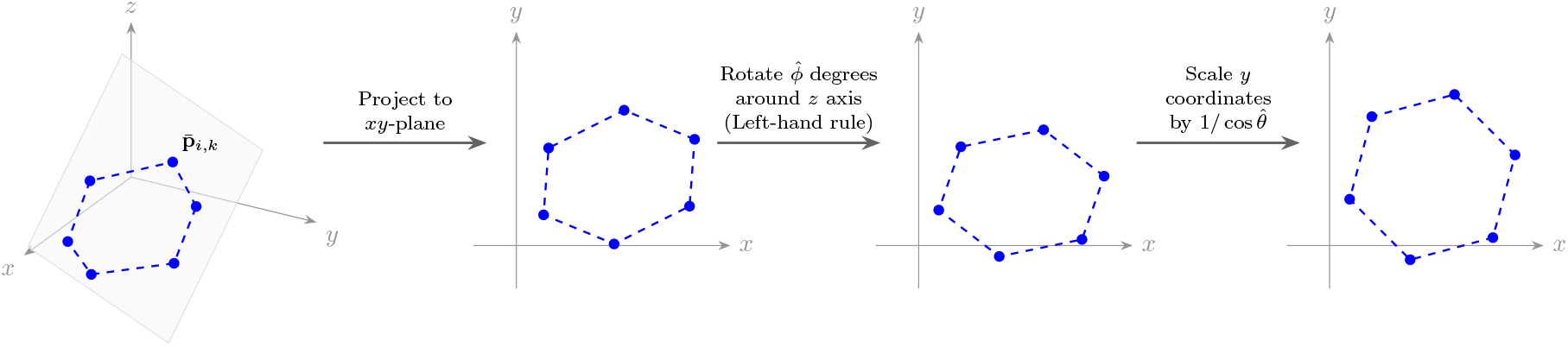
Restoration of a noise-free regular hexagonal structure from a tilted sample plane. The proposed transformations correct the geometric distortion caused by the plane’s tilt and rotation, restoring the regular hexagon’s original size.

Applying these transformations yields the 2-dimensional coordinates and the corresponding position errors,

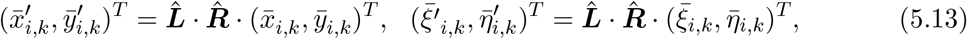

where 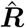 and 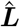 denote the two-dimensional rotation and scaling matrices, respectively, defined as:

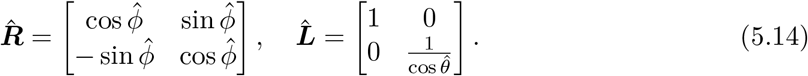

with the tilt and rotation angles 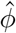 and 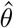 derived in the plane-fitting (Eq (5.12)).

To quantify the uncertainty of these transformed coordinates, let 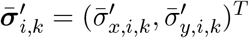denote the standard deviation vector of the transformed protomer position error 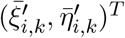. Based on the linear transformations in Eq (5.13), these standard deviation components are calculated as:

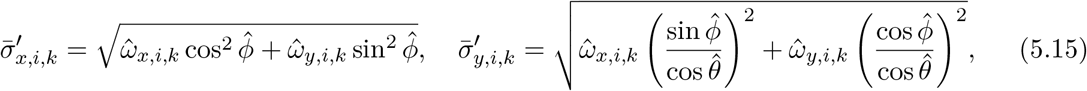

where 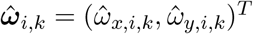 represents the variance of the original protomer position errors on the tilted sample plane introduced in Eq (5.6).

Furthermore, to bound the localization error for each oligomer *k*, we define its maximum standard deviation across all protomers and spatial dimensions as,

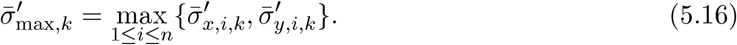

This upper bound 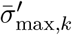is an important parameter for our subsequent analysis of the circle-fitting algorithm, and is also essential for formulating the weighted mean estimator introduced in Section 2 to downweight side length estimates with large standard deviations.

#### 5.2.2 Estimation of the oligomer side length

In this subsection, we utilize the transformed protomer positions 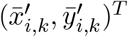and their associated localization errors 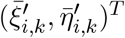 to estimate the circumcircle radius 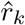for each oligomer *k*. Once 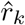 is estimated, the side length of the *k*-th oligomer can be calculated using the geometric relation:

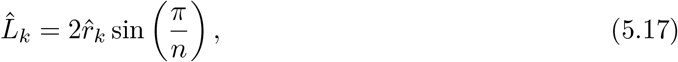

where *n* is the oligomerization.

However, estimating the circumcircle radius requires careful handling of the localization error.As a result of the coordinate transformations in Subsecection 5.2.1, the error variables 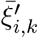 and 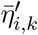are no longer independent. This correlation renders the standard geometric circle-fitting algorithm [27] used in previous work [25] not perfectly suited. To overcome this limitation, we employ the algebraic circle-fitting method.

Algebraic circle-fitting methods are widely used in practice. Unlike the geometric circle-fitting method, which iteratively minimizes the non-linear distance from the data points to the circle, algebraic circle-fitting methods minimize the sum of squared algebraic distances. This approach transforms the complex non-linear optimization problem into a fast, closed-form quadratic problem. Taking our case for example, for each oligomer *k*, one has to minimize the following algebraic distance:

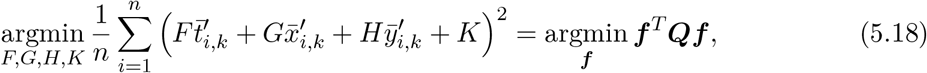

Where 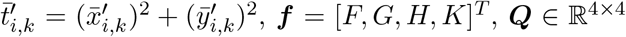 is defined as 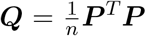, and the data matrix ***P*** ∈ ℝ^*n*×4^ is defined as,

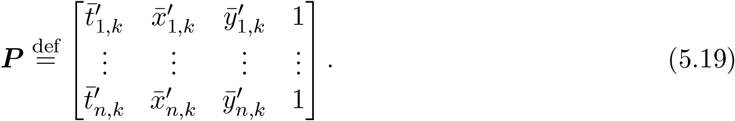

The relation between the estimator ***f*** = [*F, G, H, K*]^*T*^ and the circumcircle parameters (center (*a, b*) and radius *r*) is given by:

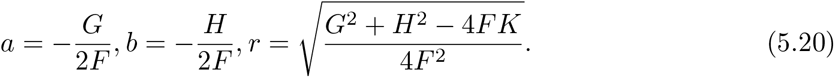

All algebraic circle-fitting methods minimize the same objective function (5.18), subject to a constraint of the form ***f*** ^*T*^ ***Uf*** = 1 to avoid trivial solution ***f*** = **0**. By incorporating the constraint using the Lagrange multiplier, we derive the matrix form of the objective function:

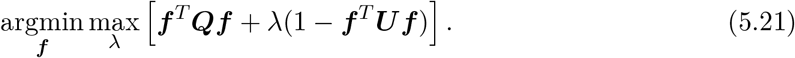

In solving min-max problem (5.21), different choices of the constraint matrix ***U*** distinguish methods such as the Kåsa [10, 6, 5], Pratt [18], Taubin [32], and hyperaccurate fits [5], ultimately resulting in different orders of estimator bias. A detailed summary of these constraint matrices can be found in [5]. As also introduced in [5], if the position noise of the transformed protomers 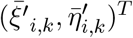is i.i.d. with zero mean and variance *σ*^2^ across all spatial directions and protomers, the bias of the estimator 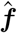derived by all algebraic circle-fitting methods can be written to,

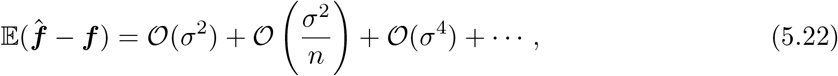

where 𝒪 (*σ*^2^) is called the essential error. By designing a specific constraint matrix ***U*** that eliminates the essential error for solving (5.21), the hyperaccurate fit is achieved. Consequently, the proposed method outperforms both standard algebraic approaches (e.g., the Kåsa, Pratt, and Taubin fits) and conventional geometric circle-fitting methods [5].

In fact, our plane-fitting method (Algorithm 1) discussed earlier is also based on this algebraic framework. Most importantly, this framework is well-suited for handling the correlated noise present in our case. Although this correlation alters the formulation of the estimator bias, the second-order error can be expressed as 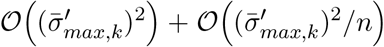. Here, the first term represents the essential error, and 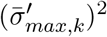is the maximum variance for each oligomer defined in (5.16).

To derive a high-precision estimator ***f*** under correlated noise, we propose constraint matrices ***U*** and 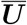, which eliminate the essential error and the entire second-order error of the estimator, respectively (see the Supporting Information 7.2 for detailed derivations). Notably, when the noise variables 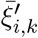 and 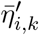 are i.i.d, our derived constraint matrices ***U*** and 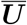 reduce to the standard constraint matrices proposed in [27] and [20], respectively. After deriving the constraint matrices, it is remained to solve the min-max problem (5.21). Parallel to both the theoretical analysis in [26] and our plane-fitting problem (5.9), the circle-fitting problem (5.21) is equivalent to finding the eigenvector 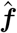 associated with the smallest positive eigenvalue *λ* of the generalized eigenvalue problem ***Qf*** = *λ****Uf*** or 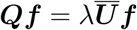.

Though we provide an explicit theoretical derivation for 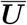 to eliminate the second-order bias, we observe that utilizing this matrix introduces numerical instability compared to using ***U***. Consequently, we employ the constraint matrix ***Ū*** in this paper to ensure stability and eliminate the essential bias for the side length estimations. On the other hand, since directly solving ***Qf*** = *λ****Uf*** for the smallest positive eigenvalue is often ill-conditioned, we stabilize it exactly as in Eq (5.11), yielding the equivalent formulation:

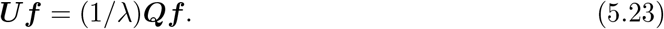

We then compute the eigenvector 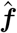 corresponding to the largest positive eigenvalue (1*/λ*) of Eq (5.23). Eventually, the complete algebraic circle-fitting method for correlated noise is summarized in Algorithm 2.

##### Algorithm 2

Hyperaccurate Algebraic Circle-Fitting Method for Correlated Noise

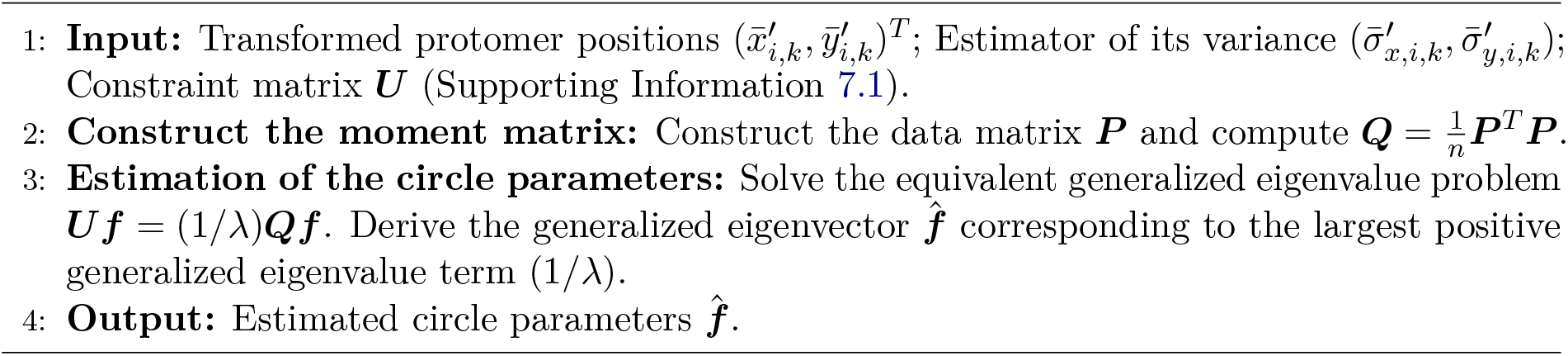

#### 5.2.3 A strategy for improving the robustness of the circle-fitting method

The circle-fitting procedure for oligomers is performed with *n* data points, where *n* is typically small (see Table 2). Consequently, the fitting algorithm is sensitive to outlier positions. To circumvent this limitation, the traditional squared-error objective is replaced with a robust penalty function that bounds the influence of large residuals [15, 8]. The robust M-estimator is then obtained by minimizing this robust objective function. Recently, [19] applied the Black-Rangarajan duality [4, 38] to reformulate this robust objective function into a weighted least-squares problem, where the weights are directly incorporated into the algebraic matrices and efficiently solved by the hyperaccurate method [27]. Motivated by this approach, we apply the M-estimation framework to our circle-fitting problem (5.18). Subsequently, by using the Black-Rangarajan duality and introducing a Lagrange multiplier to enforce the algebraic constraint, the robust optimization problem is equivalently written as the following min-max formulation:

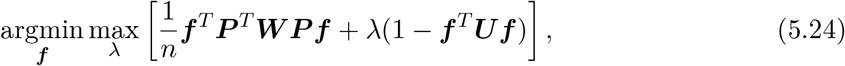

where ***P*** is the data matrix defined in (5.19), ***U*** is the constraint matrix designed for reaching a specific bias given in the Supporting Information 7.2, and ***W*** is a diagonal weight matrix derived from the Black-Rangarajan duality to enforce the M-estimation, with its explicit form given later.

To efficiently solve the robust min-max problem (5.24), we employ an iteratively reweighted least squares scheme that alternates between estimating the circle parameters and updating the weights. Specifically, at the *t*-th reweighting step, we first fix the weight matrix 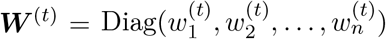. Problem (5.24) then reduces to the generalized eigenvalue problem ***P*** ^*T*^ ***W*** ^(*t*)^***P f*** = *λ****Uf***, which is solved to obtain the updated parameter vector ***f*** ^(*t*)^. Subsequently, we evaluate the current residuals 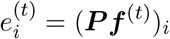. To downweight the influence of outliers for the next step, the weights are updated to 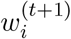 using the Huber [8] influence function, i.e.,

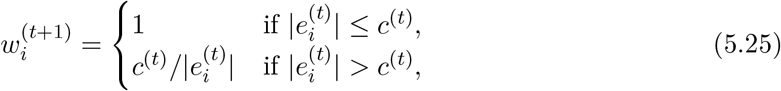

where the threshold *c*^(*t*)^ is empirically chosen as 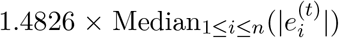. This update continues until convergence or the stopping criterion is met.

To ensure the derived estimator is free of essential bias, we design the constraint matrix ***U*** for the generalized eigenvalue problem ***P*** ^*T*^ ***W*** ^(*t*)^***P f*** = *λ****Uf*** in the Supporting Information 7.2. Compared to the work in [19], we consider correlated noise in our case, which introduces additional challenges in deriving the constraint matrix ***U***. Furthermore, this robust framework encompasses the general circle-fitting problem (5.21) as a special case of (5.24) simply by setting ***W*** = ***I***. Finally, the complete robust hyperaccurate algebraic circle-fitting for correlated noise is summarized in Algorithm 3, where the generalized eigenvalue problem is reformulated and solved for the largest positive eigenvalue term (1*/λ*) to ensure numerical stability.

##### Algorithm 3

Robust Hyperaccurate Algebraic circle-fitting Method for Correlated Noise

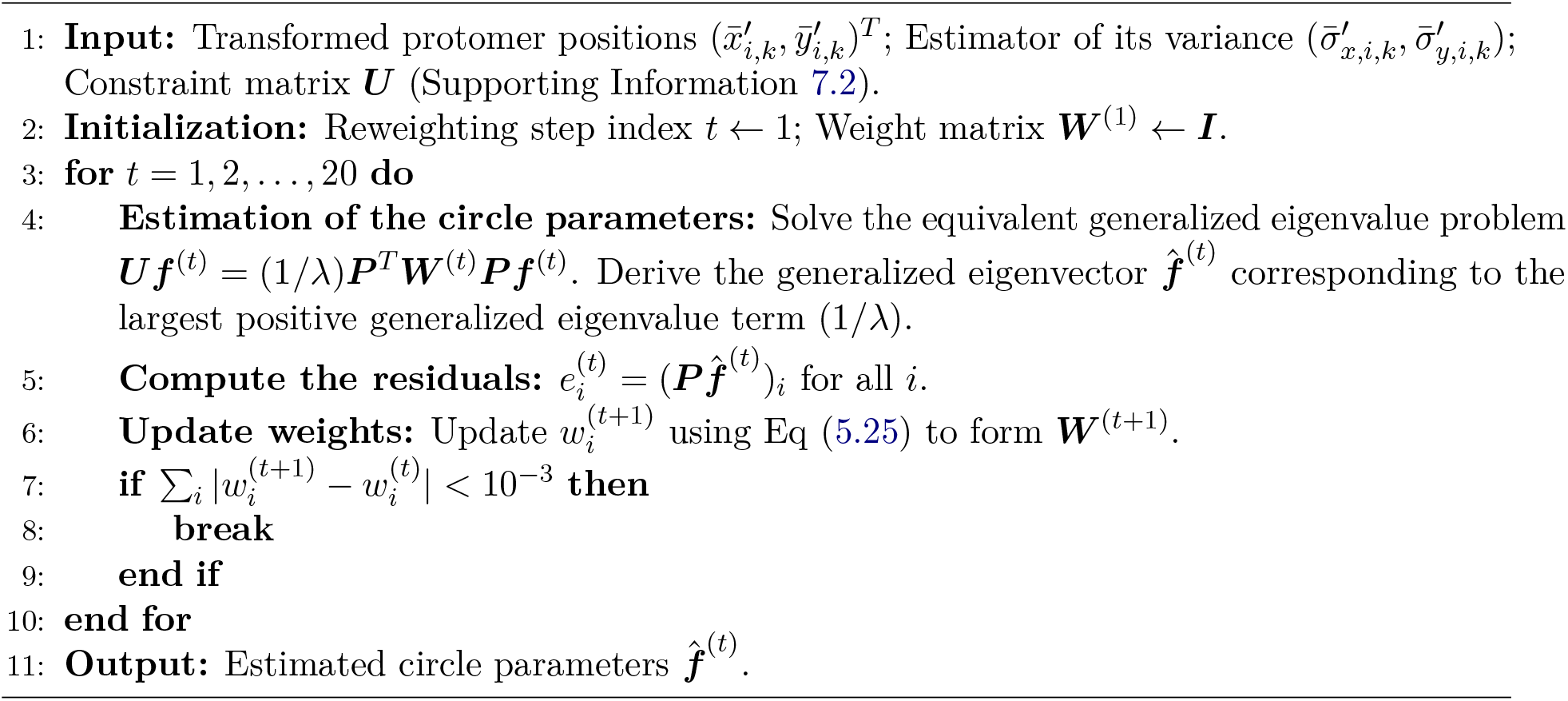

### 5.3. Postprocessing of the reconstructed side length

To ensure the reliability of the reconstructed oligomeric structures, we apply two postprocessing strategies to the results obtained from our circle-fitting algorithms (Algorithm 2, 3).

Firstly, based on the prior knowledge that the physical size of the oligomers is typically on the order of a few nanometers [1], we restrict the estimated side length to a physically meaningful range of 0 nm to 100 nm. This threshold filters out extreme and unphysical estimates while maintaining a wide enough range to test our algorithms.

Secondly, we enforce a geometric constraint on the fitted circumcircle of the oligomer. Since the oligomeric structure is modeled as a regular polygon (see Subsection 4.1), the protomer positions should be distributed across the entire fitted circle rather than clustered along a short arc, even in the presence of localization errors. To enforce this condition, we sequentially evaluate the positive eigenvalues of the generalized eigenvalue problems in Algorithms 2 and 3 in descending order, selecting the first one whose corresponding eigenvector yields a valid center. If none of the positive eigenvalues satisfy this condition, the oligomer is considered invalid and is discarded.

## 6. Summary of the workflow

To summarize, by integrating data preprocessing, the plane-fitting method, geometric transformations, the circle-fitting algorithms, and postprocessing strategies, we establish two comprehensive workflows for determining oligomeric structures: the Nominal workflow and the Robust workflow. The difference between them lies in the choice of the circle-fitting algorithm: the Nominal workflow employs circle fiting method (Algorithm 2), whereas the Robust workflow employs the robust circle-fitting method (Algorithm 3). Table 4 summarizes these workflows, which are designed for unknown sample planes with arbitrary tilt or rotation relative to the focal plane.

## Acknowledgements

This research was funded in whole, or in part, by the Austrian Science Fund (FWF) 10.55776/P34981 (OS & YD & ZY) – New Inverse Problems of Super-Resolved Microscopy (NIPSUM), SFB 10.55776/F68 (OS) “Tomography Across the Scales,” project F6807-N36 (Tomography with Uncertainties). The financial support by the Austrian Federal Ministry for Digital and Economic Affairs, the National Foundation for Research, Technology and Development and the Christian Doppler Research Association is gratefully acknowledged.MCS was funded by Howard Hughes Medical Institute, USA, Janelia Research Campus. GJS was funded by project F6809-N36 (Ultra-high Resolution Microscopy). For the purpose of open access, the author has applied a CC BY public copyright license to any Author Accepted Manuscript version arising from this submission.

## Author Contributions

**Conceptualization:** Yakun Dong, Zheyi Yang, Magdalena C. Schneider, Otmar Scherzer, Gerhard J. Schütz.

**Formal analysis:** Yakun Dong, Zheyi Yang.

**Funding acquisition:** Otmar Scherzer, Gerhard J. Schütz.

**Investigation:** Yakun Dong, Zheyi Yang, Magdalena C. Schneider.

**Methodology:** Yakun Dong, Zheyi Yang, Magdalena C. Schneider, Otmar Scherzer, Gerhard J. Schütz.

**Resources:** Otmar Scherzer, Gerhard J. Schütz.

**Software:** Yakun Dong, Zheyi Yang, Magdalena C. Schneider.

**Supervision:** Otmar Scherzer, Gerhard J. Schütz.

**Writing:** Yakun Dong, Zheyi Yang, Magdalena C. Schneider, Otmar Scherzer, Gerhard J. Schütz.

## 7. Supporting information

In this section, we introduce the error analysis for designing the constraint matrices for the plane-fitting problem (5.9) and the robust circle-fitting problem (5.24), alongside a comparison of our estimation results with the prior work in [25]. Given that both problems are based on the same algebraic fitting framework, we provide a detailed analysis of the plane-fitting problem and a more concise analysis of the robust circle-fitting problem (5.24).

### 7.1. Error analysis for designing the constraint matrix *N* for plane-fitting problem (5.9)

We analyze the plane-fitting problem (5.9) in detail to design the corresponding constraint matrix. Specifically, the fitting is performed based on all estimated protomer positions 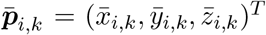, with the position errors 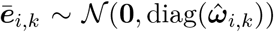,where 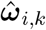is defined in (5.6). To evaluate the impact of these errors rigorously, we apply the matrix perturbation method [31] within a high-order error analysis framework [9, 27].

Recall the min-max plane-fitting problem (5.9),

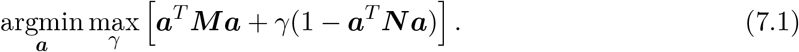

The optimal parameter vector âis obtained when ***a*** and *γ* satisfy the optimality conditions: ***Ma*** = *γ****Na*** and ***a***^*T*^ ***Na***− 1 = 0. By multiplying both sides of the first condition by ***a***^*T*^ and substituting the constraint, we derive

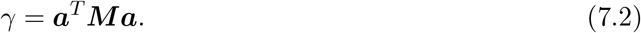

Thus, the problem (5.9) is equivalent to finding the smallest positive generalized eigenvalue *γ* and its corresponding generalized eigenvector ***â*** by solving the equation

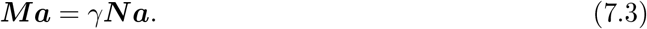

We then define the maximum standard deviation across all protomers and oligomers along all spatial directions as,

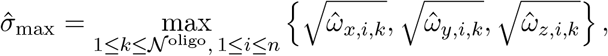

Where 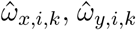 and 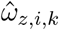 are the components of 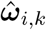, defined in (5.6). Then we use the matrix perturbation method and expand all data-dependent quantities (***â***, ***M***, and ***N***) in terms of the noise level as,

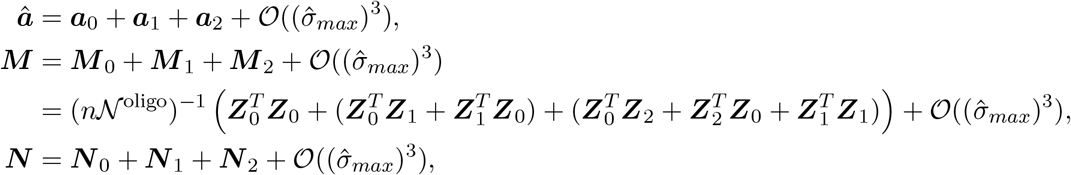

Where

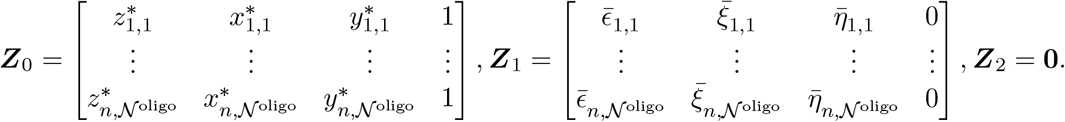

with 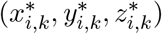being the ground-truth protomer position, and 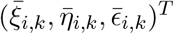being the protomer position error defined in (5.3). The variables with the subscript (*·*)_0_ represent the groundtruth quantities (zero-th order perturbation terms), while those with (*·*)_1_ and (*·*)_2_ represent the first-order and the second-order perturbation terms, respectively. The remainder term 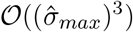represents a random variable ℛ such that 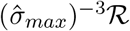 is bounded in probability.

With these preparations, we first analyze the variance of the plane estimator â. Since the ground-truth data points lie on the ground-truth tilted sample plane, it follows that

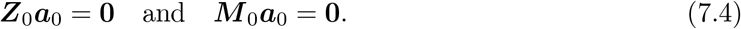

Thus, we can analyze the perturbation behavior of the Lagrange parameter *γ* in (7.1) as follows:

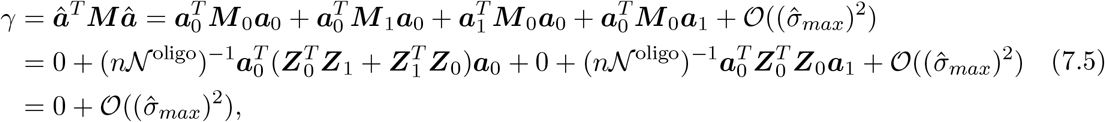

where the first equality is from relation (7.2) and the third and forth equality is based on relation (7.4). As shown in Eq (7.5), both the zero-th and first-order terms in the perturbation expansion of *γ* vanish, indicating that *γ* is a second-order quantity with respect to the noise level, i.e. 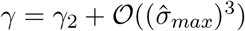.

Next, we substitute the perturbation expansions into the generalized eigenvalue problem ***Ma*** = *γ****Na*** (7.3) and match terms of the same order. Since *γ* lacks zeroth- and first-order terms, these components vanish completely on the right-hand side of expansions of (7.3). Thus, setting the zeroth- and first-order terms of ***Ma*** to zero yields:

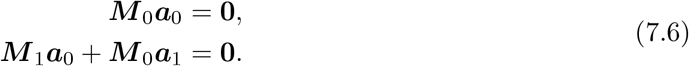

Subsequently,

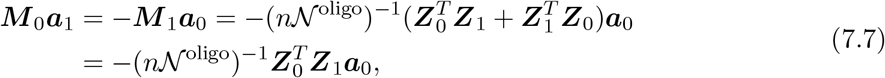

where we expand ***M*** _1_ as in (7.1) and use the relations (7.4). Then we derive the formula for ***a***_1_ as follows:

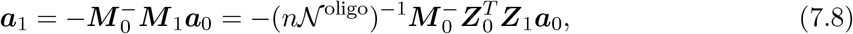

Where 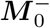is the Moore–Penrose inverse. Thus, the first order bias of the plane estimator âis

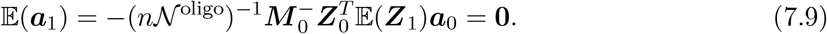

We also observe that ***a***_1_ is independent of ***N***. Consequently, the dominant term of the estimator variance is given by

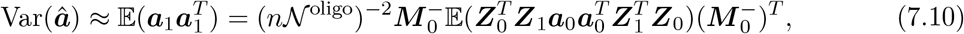

which is also independent of ***N***. Next, we examine the bias behavior with respect to different ***N***. Again, we expand both sides of the eigenvalue problem ***Ma*** = *γ****Na*** (7.3), and match the second order of the errors,

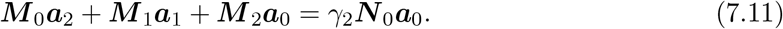

By multiplying 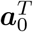 on both sides of Eq (7.11) and substituting ***a***_1_ by Eq (7.8), we derive the expression of *γ*_2_ as

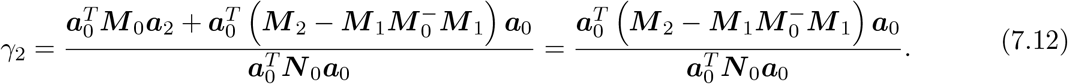

Substituting the expression of *γ*_2_ into Eq (7.11), we derive ***a***_2_ as

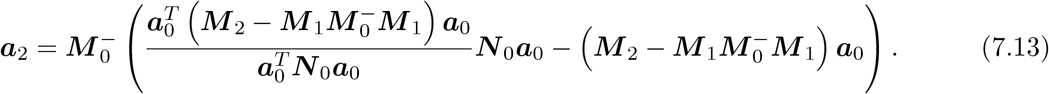

Thus, the bias to the second order is given by

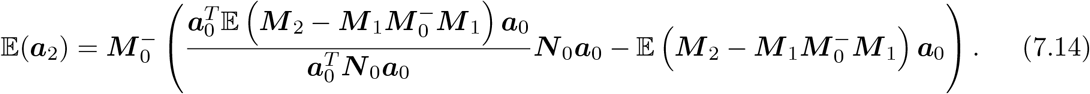

Therefore, combining (7.9) and (7.10), we observe that regardless of the constraint matrix ***N***, the estimator ***â*** remains first-order unbiased with an identical dominant variance. However, its second-order bias strongly depends on ***N***_0_. In particular, if we select

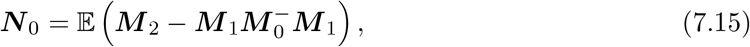

then 𝔼 (***a***_2_) = 0, which means the estimator ***â*** is unbiased up to second order in the error, where ***M*** _0_ is the noise-free data term. Then, in the next subsection, we base on ***N*** _0_ (7.15) to construct the constraint matrix ***N*** by adding noise terms.

#### Expression of a possible constraint matrix *N* for plane-fitting

In this subsection, we provide a detailed form of the constraint matrix ***N*** for the plane-fitting problem. Based on ***N*** _0_, which is the zeroth-order term in the perturbation of the constraint matrix ***N***, various realizations of ***N*** can be constructed. Here, we present one possible choice.

First, we expand the expression for ***N*** _0_:

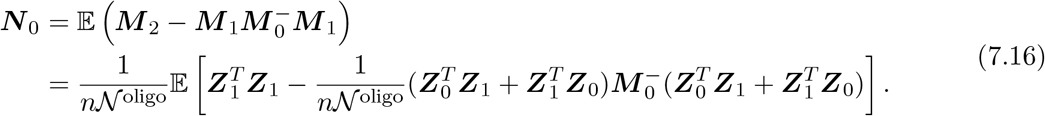

Define a vector 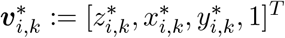 for *i* ∈ {1, …, *n*} and *k* ∈ {1, …, 𝒩 ^oligo^}, and matrices 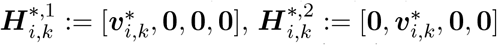 and 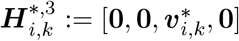, we can derive the different components for ***N*** _0_ in Eq (7.16) as follows:

- 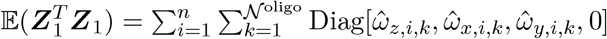
- Expression of 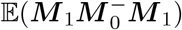:

Since 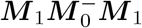 can be split into 4 terms,

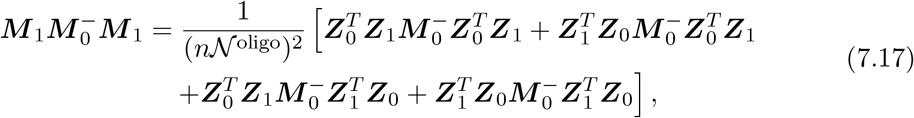

we showcase the derivation for 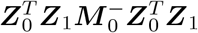; the other terms can be treated analogously. Define a matrix sequence 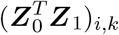 with index *i* and *k* as

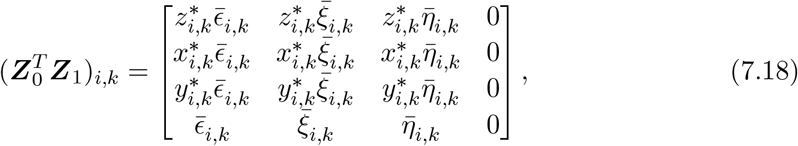

Thus 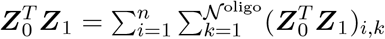. Furthermore, by the definition of 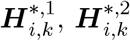 and 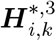 we have

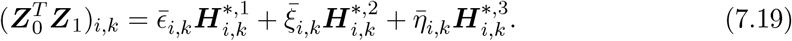

Since 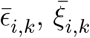 and 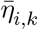 are independent and identically distributed random variables, the expectation of 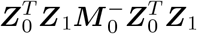 can be written as

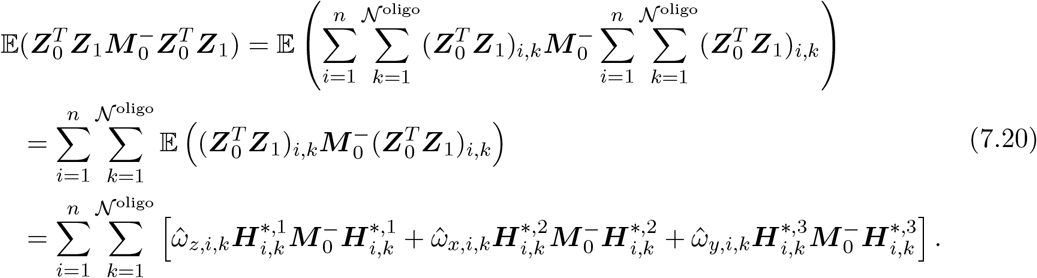

Thus,

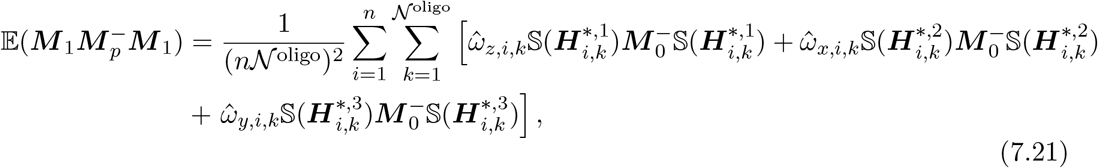

where 𝕊 (*·*) = ((*·*) + (*·*)^*T*^) is the symmetrization operator.

In summary, the expression of ***N*** _0_ is given as

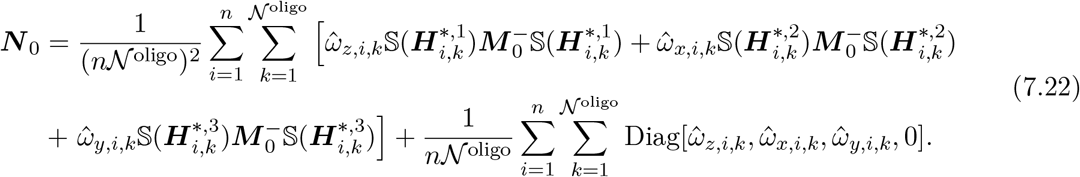

Then, by plugging the estimates of 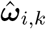defined in (5.6) into (5.6), and replacing the ground-truth vector 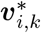in 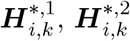 and 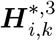 by noisy 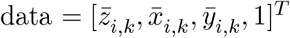, we derive one possible ***N*** for the plane-fitting problem (5.9) that ensures the estimator ***â*** remains bias-free up to second order in the error.

Since we typically simulate a lot of oligomers in the sample plane, the term 1*/*(*n*𝒩 ^oligo^) will dominate over 1*/*(*n*𝒩 ^oligo^)^2^. By neglecting the term 1*/*(*n*𝒩 ^oligo^)^2^ in Eq (7.22) and substituting the estimates 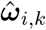from (5.6), we derive one simplified possible ***N*** for the plane-fitting problem (5.9) as

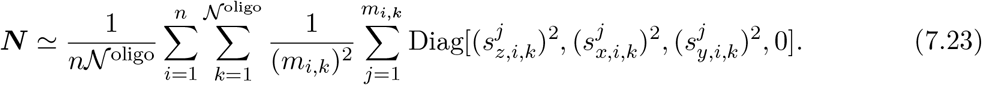

Finally, we derive a constraint matrix (7.22), together with a simplified one in (7.23), which eliminates the second-order error of the plane estimator ***â***.

### 7.2. Error analysis for designing the constraint matrix *U* for (robust) circle-fitting problem (5.24)

Analogous to the plane-fitting analysis in subsection 7.1, we now evaluate the robust circle-fitting problem (5.24). Specifically, the fitting is performed based on the transformed protomer positions 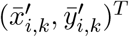, with the transformed position errors 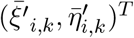 defined in Subsection 5.2.1.

Since the standard circle-fitting problem (5.21) follows as a special case with ***W*** = ***I***, we directly analyze the robust formulation. Specifically, we recall the robust circle-fitting problem (5.24):

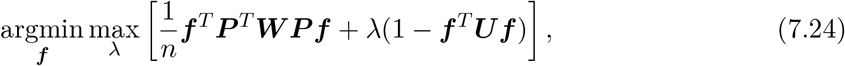

where ***P*** is the data matrix defined in Eq (5.19), ***U*** is the constraint matrix to be designed later, and ***W*** is a diagonal weight matrix. By analyzing the optimality condition of problem (7.24), solving this min-max problem is equivalent to solving the generalized eigenvalue problem ***P*** ^*T*^ ***W P f*** = *λ* ***Uf***.

Then we use the matrix perturbation method and expand all data-dependent quantities (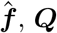, ***Q***, and ***U***) in terms of the noise level as,

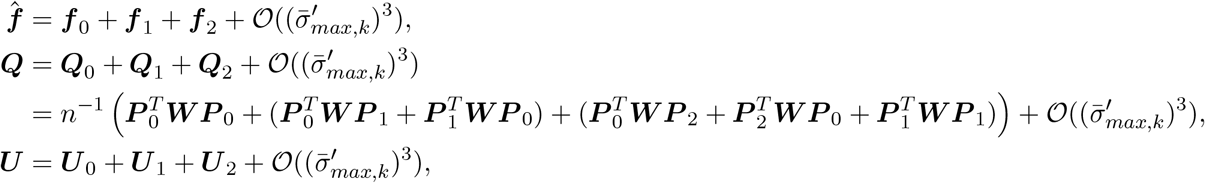

where 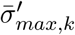is the maximum standard deviation across all protomers and spatial dimensions in one transformed oligomer, defined in (5.16). The expansion of the data matrix ***P*** is given as,

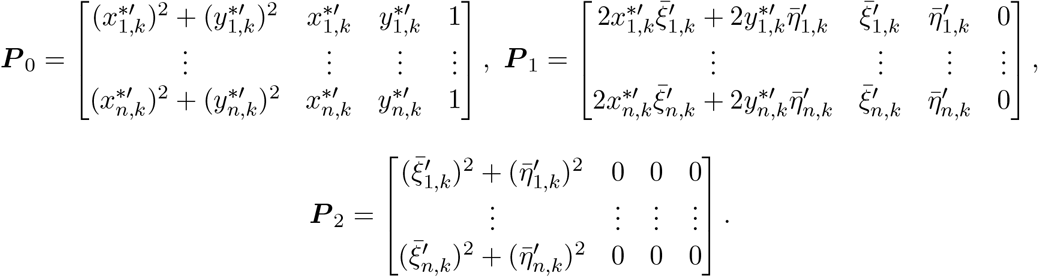

where 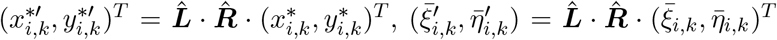, with 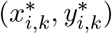 being the ground-truth *x*-*y* direction protomer position and 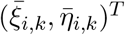being the *x*- and *y*-direction protomer position error defined in Eq (5.3) and 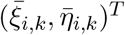being the transformation matrix defined in Subsubsection 5.2.1.

Following a perturbation procedure similar to that in Supporting Information 7.1, we analyze the optimality condition of problem (5.24), namely ***P*** ^*T*^ ***W P f*** = *λ* ***Uf***. This yields the first-order expansion of the circle parameter 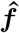as follows:

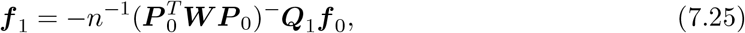

where [·]^−^ is the Moore-Penrose pseudoinverse of the matrix [·]. From Eq (7.25), we can observe that ***f*** _1_ is independent of the constraint matrix ***U***, and the first-order bias of the estimator is zero, i.e., 𝔼 (***f*** _1_). Furthermore, the dominant term of the estimator variance is given by

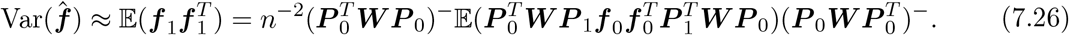

which also concludes that the leading term of the variance of the circle estimator 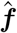is independent of the choice of the constraint matrix ***U***.

Next, we examine the second-order bias of the circle estimator 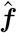. By setting this bias to zero, we obtain the constraint matrix

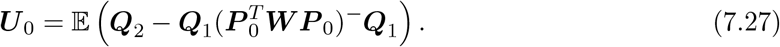

Based on the ***U*** _0_, which is the zeroth-order term in the perturbation of the constraint matrix ***N***, we can construct a constraint matrix 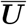 by replacing the ground-truth terms with their corresponding noisy observations. By using this constraint matrix in the circle-fitting problem, we obtain an estimator 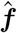 that is free of second-order bias.

However, in numerical experiments, we observe instability when solving the eigenvalue problem using ***Ū*** to remove the entire second-order bias of 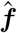. To improve robustness, we omit the term 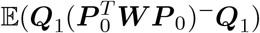,whose magnitude is 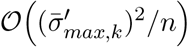. Instead, we only keep the term 𝔼 (***Q***_2_) with a magnitude of 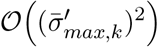, which is essential bias of the estimator. Using this simplified constraint matrix ***U***, we obtain the hyperaccurate algebraic circle-fitting algorithms under correlated noise (Algorithms 2 and 3). Additionally, when the error variables 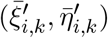 are independent and share the same standard deviation across all spatial directions, our method naturally reduces to the algorithm introduced in [20] when using the constraint matrix 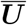, and to the method in [27] when using ***U***.

#### Possible constraint *U* for Algorithm 2 and 3

Expand the constraint matrix ***U*** _0_ = 𝔼E(***Q***_2_),we have,

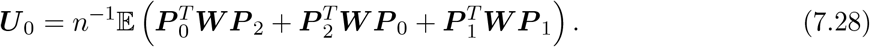

Note that we have two more terms compared to the expansion in Eq (7.16), which are 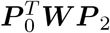, and 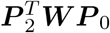. This is due to the second-order expansion of the data term, i.e., ***P*** _2_ being non-zero in this case.

Define a vector 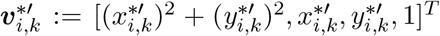 for *i* ∈ {1, …, *n*}, and matrix 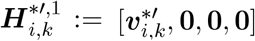, where 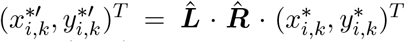. With these, we can derive different components for ***U*** _0_ in Eq (7.28) as follows:

- 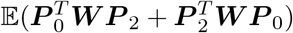 is given as

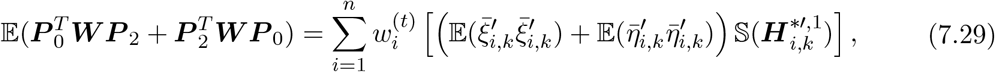
- 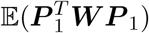 is given as

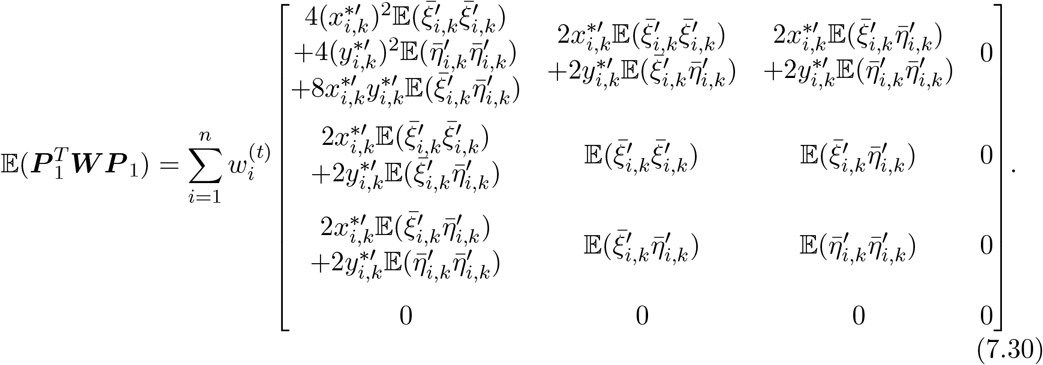

The expectation of the noise terms are given as

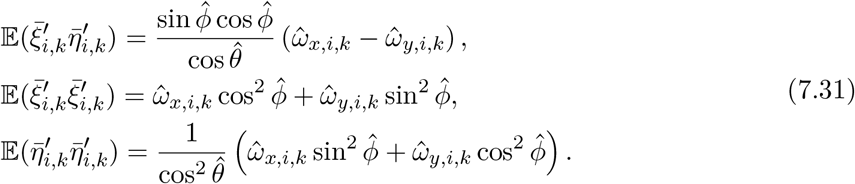

where the calculation is based on the transformation relation (5.13), with 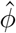and 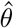representing the estimated tilt and rotation angles of the sample plane given in (5.12), and 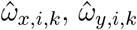, and 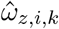 being the components of 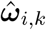 defined in (5.6).

To derive the constraint matrix ***U*** that eliminate the essential bias of the circle estimator 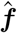from (7.28), we first plug the estimates 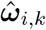from (5.6) into relation (7.31), and substitute the expectations in (7.28). Next, we replace the ground-truth term 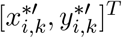 in 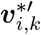 and 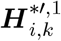 with the noisy data 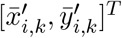 in (7.28). Through these steps, we derive a possible matrix ***U*** for the circle-fitting problem (5.21) and the robust circle-fitting problem (5.24).

### 7.3. Comparison results with previous methods

In this part, we compare our proposed Robust workflow and the workflow in [25]. However, the workflow in [25] is not directly applicable to our setting, because the tilted sample plane is unknown, and to the author’s knowledge, the geometric circle-fitting method used in [25] cannot be extended to cases with anisotropic localization precision in the *x* and *y* directions. For the purpose of comparison, we approximate the localization precision of each protomer by averaging the precisions in the *x* and *y* directions, and then apply the geometric circle-fitting method directly to the *x* and *y* positions of the protomers.

The simulation setting follows that of Simulation I in the main text: we simulate 5000 tetramers with a fixed side length *L*^*\**^ = 5 nm on sample planes tilted and rotated by 0°, 10°, 20°, and 30°. The photon count is set to *N*_max_ = 5 ×10^6^ for all simulations. In our workflows, we use the weighted mean estimator to select representative estimates, whereas for the workflow in [25], we use the median estimator, which is considered efficient for geometric circle-fitting in their study. The comparison results are summarized in Table 5. As shown, when the sample plane is perfectly aligned with the focal plane, the workflow in [25] performs slightly better. This is due to slight anisotropies in the localization precision, and the fact that the geometric circle-fitting method extended in [25] eliminates the entire second-order bias, while our algebraic fitting method only eliminates the essential error (introduced in Subsection 5.2.2). However, when the sample plane is tilted relative to the focal plane, our workflow becomes necessary for high-precision estimations.

**Table 5.** Comparison between our proposed workflows (with plane-fitting) and the original workflow in [25]. The first column shows the ground-truth tilt and rotation of the simulated sample plane. Columns 3 and 4 show the tilt and rotation angles of the fitted plane, and the chosen estimator for each workflow.

| $\theta/\phi$ ($^\circ$) | Workflow | $\hat{\theta}/\hat{\phi}$ ($^\circ$) | Chosen estimator for each workflow (nm) |
| --- | --- | --- | --- |
| 0/0 | Robust workflow<br>Workflow in [25] | 0.0001 / -<br>- | 5.0017<br>**4.9994** |
| 10/10 | Robust workflow<br>Workflow in [25] | 10.0000 / 10.0002<br>- | **5.0024**<br>4.9602 |
| 20/20 | Robust workflow<br>Workflow in [25] | 20.0000 / 20.0001<br>- | **5.0037**<br>4.8476 |
| 30/30 | Robust workflow<br>Workflow in [25] | 30.0000 / 30.0000<br>- | **5.0064**<br>4.6676 |

### 7.4. Software

The latest version of the software will be uploaded upon submission to the journal.

## References

[1] B. Alberts, A. Johnson, J. Lewis, M. Raff, K. Roberts, and P. Walter. “Molecular Biology of the Cell”. Garland Science, 2014 (cited on page 17).

[2] A. M. Arnold, M. C. Schneider, C. Hüsson, et al. “Verifying molecular clusters by 2-color localization microscopy and significance testing”. In: Scientific Reports 10 (2020), p. 4230. DOI: 10.1038/s41598-020-60976-6 (cited on page 8).

[3] P. R. Bevington and D. K. Robinson. “Data reduction and error analysis for the physical sciences”. 3rd. New York: McGraw-Hill, 2003 (cited on page 4).

[4] M. J. Black and A. Rangarajan. “On the unification of line processes, outlier rejection, and robust statistics with applications in early vision”. In: International journal of computer vision 19.1 (1996), pp. 57–91 (cited on page 15).

[5] N. Chernov and C. Lesort. “Least Squares Fitting of Circles”. In: Journal of Mathematical Imaging and Vision 23.3 (2005), pp. 239–252. ISSN: 1573-7683. DOI: 10.1007/s10851-005-0482-8 (cited on pages 14, 15).

[6] P. Delogne. “Computer optimization of Deschamps’ method and error cancellation in reflectometry”. In: Proc. IMEKO-Symp. Microwave Measurements. 1972, pp. 117–123 (cited on page 14).

[7] F. Hinterer, M. C. Schneider, S. Hubmer, M. López-Martinez, P. Zelger, A. Jesacher,R. Ramlau, and G. J. Schütz. “Robust and bias-free localization of individual fixed dipole emitters achieving the Cramér Rao bound for applications in cryo-single molecule localization microscopy”. In: PLoS ONE 17.2 (2022). Ed. by M. Cebecauer, e0263500. DOI: 10.1371/journal.pone.0263500 (cited on pages 2, 8, 9).

[8] P. J. Huber. “Robust Estimation of a Location Parameter”. In: The Annals of Mathematical Statistics 35.1 (1964), pp. 73–101. DOI: 10.1214/aoms/1177703732 (cited on pages 15, 16).

[9] K. Kanatani. “Statistical optimization for geometric fitting: Theoretical accuracy bound and high order error analysis”. In: International Journal of Computer Vision 80.2 (2008), pp. 167–188 (cited on pages 11, 19).

[10] I. Kasa. “A curve fitting procedure and its error analysis”. In: IEEE Trans. Instrum. Meas. IM-25.1 (1976), pp. 8–14 (cited on page 14).

[11] R. Kaufmann, C. Hagen, and K. Grünewald. “Fluorescence cryo-microscopy: current challenges and prospects”. In: Current Opinion in Chemical Biology 20 (2014), pp. 86–91. ISSN: 1367-5931. doi: 10.1016/j.cbpa.2014.05.007 (cited on page 1).

[12] M. D. Lew, M. P. Backlund, and W. E. Moerner. “Rotational Mobility of Single Molecules Affects Localization Accuracy in Super-Resolution Fluorescence Microscopy”. In: Nano Letters 13.9 (2013), pp. 3967–3972. DOI: 10.1021/nl304359p (cited on page 2).

[13] W. Li, S. C. Stein, I. Gregor, and J. Enderlein. “Ultra-stable and versatile widefield cryo-fluorescence microscope for single-molecule localization with sub-nanometer accuracy”. In: Opt. Express 23.3 (Feb. 2015), pp. 3770–3783. DOI: 10.1364/OE.23.003770 (cited on page 1).

[14] Y. Li et al. “The effects of chemical fixation on the cellular nanostructure”. In: Experimental Cell Research 358.2 (2017), pp. 253–259. ISSN: 0014-4827. DOI: 10.1016/j.yexcr.2017.06.022 (cited on page 1).

[15] R. A. Maronna, R. D. Martin, V. J. Yohai, and M. Salibián-Barrera. “Robust statistics: theory and methods (with R)”. John Wiley & Sons, 2019 (cited on page 15).

[16] H. Mazal, F.-F. Wieser, D. Bollschweiler, A. Schambony, and V. Sandoghdar. “Cryo–light microscopy with angstrom precision deciphers structural conformations of PIEZO1 in its native state”. In: Science Advances 11.34 (2025), eadw4402. DOI: 10.1126/sciadv.adw4402. eprint: https://www.science.org/doi/pdf/10.1126/sciadv.adw4402 (cited on page 2).

[17] H. Mazal, F.-F. Wieser, and V. Sandoghdar. “Deciphering a hexameric protein complex with Angstrom optical resolution”. In: eLife 11 (May 2022). Ed. by F. Campelo, V. Dötsch, and S. Brasselet, e76308. ISSN: 2050-084X. DOI: 10.7554/eLife.76308 (cited on page 2).

[18] V. Pratt. “Direct least-squares fitting of algebraic surfaces”. In: SIGGRAPH Comput. Graph. 21.4 (Aug. 1987), pp. 145–152. ISSN: 0097-8930. DOI: 10.1145/37402.37420 (cited on page 14).

[19] Z. Qi, W. Wang, T. Luo, W. Cheng, and Z. Liu. “A robust circle fitting method for component fiducialization”. In: Nuclear Instruments and Methods in Physics Research Section A: Accelerators, Spectrometers, Detectors and Associated Equipment 1068 (2024), p. 169775. ISSN: 0168-9002. DOI: 10.1016/j.nima.2024.169775 (cited on pages 15, 16).

[20] P. Rangarajan and K. Kanatani. “Improved algebraic methods for circle fitting”. In: Electronic Journal of Statistics 3.none (2009), pp. 1075–1082. DOI: 10.1214/09-EJS488 (cited on pages 15, 24).

[21] B. Rieger and S. Stallinga. “The Lateral and Axial Localization Uncertainty in Super-Resolution Light Microscopy”. In: ChemPhysChem 15.4 (2014), pp. 664–670. DOI: 10.1002/cphc.201300711. eprint: https://chemistry-europe.onlinelibrary.wiley.com/doi/pdf/10.1002/cphc.201300711 (cited on page 2).

[22] B. Rossboth, A. M. Arnold, H. Ta, R. Platzer, F. Kellner, J. B. Huppa, M. Brameshuber, F. Baumgart, and G. J. Schütz. “TCRs Are Randomly Distributed on the Plasma Membrane of Resting Antigen-Experienced T Cells”. In: Nature Immunology 19.8 (Aug. 2018), pp. 821–827. DOI: 10.1038/s41590-018-0162-7 (cited on page 8).

[23] L. Schermelleh, A. Ferrand, T. Huser, C. Eggeling, M. Sauer, O. Biehlmaier, and G. P. C. Drummen. “Super-Resolution Microscopy Demystified”. In: Nature Cell Biology 21.1 (Jan. 2019), pp. 72–84. DOI: 10.1038/s41556-018-0251-8 (cited on page 1).

[24] M. C. Schneider, F. Hinterer, A. Jesacher, and G. J. Schütz. “Interactive simulation and visualization of point spread functions in single molecule imaging”. In: Optics Communications 560 (2024), p. 130463. ISSN: 0030-4018. DOI: 10.1016/j.optcom.2024.130463 (cited on pages 3, 9).

[25] M. C. Schneider, R. Telschow, G. Mercier, M. López-Martinez, O. Scherzer, and G. J. Schütz. “A workflow for sizing oligomeric biomolecules based on cryo single molecule localization microscopy”. In: PLOS ONE 16 (Jan. 2021), pp. 1–23. DOI: 10.1371/journal.pone.0245693 (cited on pages 2–4, 7–9, 14, 19, 25, 26).

[26] A. Al-Sharadqah. “Further statistical analysis of circle fitting”. In: Electronic Journal of Statistics 8.2 (2014), pp. 2741–2778. DOI: 10.1214/14-EJS971 (cited on page 15).

[27] A. Al-Sharadqah and N. Chernov. “Error analysis for circle fitting algorithms”. In: Electronic Journal of Statistics 3.none (2009), pp. 886–911. DOI: 10.1214/09-EJS419 (cited on pages 11, 14, 15, 19, 24).

[28] Y. M. Sigal, R. Zhou, and X. Zhuang. “Visualizing and discovering cellular structures with super-resolution microscopy”. In: Science 361.6405 (2018), pp. 880–887. doi: 10.1126/science.aau1044. eprint: https://www.science.org/doi/pdf/10.1126/science.aau1044 (cited on page 1).

[29] C. S. Smith, N. Joseph, B. Rieger, and K. A. Lidke. “Fast, single-molecule localization that achieves theoretically minimum uncertainty”. In: Nature Methods 7.5 (May 2010), pp. 373–375. DOI: 10.1038/nmeth.1449 (cited on page 2).

[30] S. Stallinga and B. Rieger. “Accuracy of the Gaussian Point Spread Function model in 2D localization microscopy”. In: Opt. Express 18.24 (Nov. 2010), pp. 24461–24476. DOI: 10.1364/OE.18.024461 (cited on page 2).

[31] G. W. Stewart and J.-g. Sun. “Matrix Perturbation Theory”. Academic press, 1990 (cited on pages 11, 19).

[32] G. Taubin et al. “Estimation of planar curves, surfaces, and nonplanar space curves defined by implicit equations with applications to edge and range image segmentation”. In: IEEE transactions on pattern analysis and machine intelligence 13.11 (1991), pp. 1115–1138 (cited on page 14).

[33] T. K. Tsang, E. A. Bushong, D. Boassa, J. Hu, B. Romoli, S. Phan, D. Dulcis, C.-Y. Su, and M. H. Ellisman. “High-quality ultrastructural preservation using cryofixation for 3D electron microscopy of genetically labeled tissues”. In: eLife 7 (May 2018). Ed. by M. Helmstaedter, e35524. ISSN: 2050-084X. DOI: 10.7554/eLife.35524 (cited on pages 1, 9).

[34] M. W. Tuijtel, A. J. Koster, S. Jakobs, F. G. A. Faas, and T. H. Sharp. “Correlative Cryo Super-Resolution Light and Electron Microscopy on Mammalian Cells Using Fluorescent Proteins”. In: Scientific Reports 9.1 (Feb. 2019), p. 1369. DOI: 10.1038/s41598-018-37728-8 (cited on page 1).

[35] S. Weisenburger, D. Boening, B. Schomburg, K. Giller, S. Becker, C. Griesinger, and V. Sandoghdar. “Cryogenic Optical Localization Provides 3D Protein Structure Data with Angstrom Resolution”. In: Nature Methods 14.2 (Feb. 2017), pp. 141–144. DOI: 10.1038/nmeth.4141 (cited on pages 1, 9).

[36] S. Weisenburger, D. Boening, B. Schomburg, K. Giller, S. Becker, C. Griesinger, and V. Sandoghdar. “Cryogenic optical localization provides 3D protein structure data with Angstrom resolution”. In: Nature Methods 14.2 (2017), pp. 141–144. DOI: 10.1038/nmeth.4141 (cited on page 2).

[37] S. Weisenburger, B. Jing, D. Hänni, L. Reymond, B. Schuler, A. Renn, and V. Sandoghdar. “Cryogenic Colocalization Microscopy for Nanometer-Distance Measurements”. In: ChemPhysChem 15.4 (2014), pp. 763–770. DOI: 10.1002/cphc.201301080 (cited on page 2).

[38] H. Yang, P. Antonante, V. Tzoumas, and L. Carlone. “Graduated non-convexity for robust spatial perception: From non-minimal solvers to global outlier rejection”. In: IEEE Robotics and Automation Letters 5.2 (2020), pp. 1127–1134 (cited on page 15).

